# Immunotherapy-related cognitive impairment after CAR T cell therapy in mice

**DOI:** 10.1101/2024.05.14.594163

**Authors:** Anna C. Geraghty, Lehi Acosta-Alvarez, Maria Rotiroti, Selena Dutton, Michael R. O’Dea, Pamelyn J. Woo, Haojun Xu, Kiarash Shamardani, Rebecca Mancusi, Lijun Ni, Sara B. Mulinyawe, Won Ju Kim, Shane A. Liddelow, Robbie G. Majzner, Michelle Monje

## Abstract

Persistent central nervous system (CNS) immune dysregulation and consequent dysfunction of multiple neural cell types is central to the neurobiological underpinnings of a cognitive impairment syndrome that can occur following traditional cancer therapies or certain infections. Immunotherapies have revolutionized cancer care for many tumor types, but the potential long-term cognitive sequelae are incompletely understood. Here, we demonstrate in mouse models that chimeric antigen receptor (CAR) T cell therapy for both CNS and non-CNS cancers can impair cognitive function and induce a persistent CNS immune response characterized by white matter microglial reactivity and elevated cerebrospinal fluid (CSF) cytokines and chemokines. Consequently, oligodendroglial homeostasis and hippocampal neurogenesis are disrupted. Microglial depletion rescues oligodendroglial deficits and cognitive performance in a behavioral test of attention and short-term memory function. Taken together, these findings illustrate similar mechanisms underlying immunotherapy-related cognitive impairment (IRCI) and cognitive impairment following traditional cancer therapies and other immune challenges.

## Introduction

Cancer therapy-related cognitive impairment (CRCI) affects a large number of cancer survivors^1^. Traditional cancer therapies are frequently associated with a syndrome of persistent cognitive impairment characterized by deficits in memory, attention, speed of information processing, multitasking, and executive function^1,2^. While best described after traditional therapies like cytotoxic chemotherapy^1^, early reports suggest that immunotherapies may also increase the risk of persistent cognitive symptoms^3–5^ (for review see^6^). Cancer therapies such as cranial irradiation ^7–10^ or methotrexate chemotherapy^11,12,13^, can induce a reactive state in microglia, particularly in a subpopulation of microglia that reside in axon tracts^14^ such as in subcortical white matter^11^. These reactive microglia impair myelin homeostasis^11,15^ and plasticity^12^, as well as hippocampal neurogenesis^7,8^ through mechanisms that include cytokine signaling^8^ and induction of neurotoxic astrocyte substates^11,16^. Respiratory viral infections such as COVID and influenza, which can cause a cognitive syndrome that mirrors CRCI, induce a state of microglial reactivity associated with impairments in the same mechanisms of neural homeostasis and plasticity that are dysregulated after cancer therapies^15^. As microglial reactivity is central to several mechanisms underpinning cognitive impairment after traditional cancer therapies (for review, please see^17^), we hypothesized that immunotherapies - primarily inflammatory in mechanism - may similarly induce neuroinflammation, consequent dysregulation of neural cells, and cognitive impairment.

CAR T cell therapy has been transformative for refractory hematological malignancies in both children and adults, and CD19-targeting CAR T cells are now FDA-approved for treatment of acute lymphoblastic leukemias as well as other hematological cancers^18^. Recently, the disialoganglioside GD2 was found to be a promising target for a lethal high-grade glioma of childhood called H3K27M-altered diffuse midline glioma, also called diffuse intrinsic pontine glioma (DIPG) for tumors of the brainstem^19^, and early clinical trial (NCT04196413) results testing GD2-targeting CAR T cells for DIPG have been encouraging^20^. CAR T cell therapy has been associated with acute neurotoxicity syndromes such as immune effector cell-associated neurotoxicity (ICANS)^21,22^, but the longer-term consequences are understudied. To assess the neurobiological consequences of CAR T cell therapy after the acute period of tumor clearance, we tested the chronic effects of CAR T-cell immunotherapy for malignancies within and outside of the nervous system on CNS immune state, myelin homeostasis, hippocampal neurogenesis, and cognitive function in preclinical mouse models.

## Results

### Impaired cognition in mice after tumor-clearing CAR T cell therapy

To determine whether mice treated with CAR T cell therapy exhibit cognitive deficits, we utilized patient-derived xenograft murine models of CNS and non-CNS cancers (Figure 1A). These models include: an H3K27M+ diffuse intrinsic pontine glioma (DIPG) model, a pre-B cell acute lymphoblastic leukemia (ALL) model, and two osteosarcoma models. Of the two osteosarcoma models, one osteosarcoma model clears efficiently and is associated with minimal cytokine release with CAR T cell therapy (henceforth referred to as “rapid-clearing osteosarcoma”), and one osteosarcoma model is more aggressive, is engineered to express CD19 for experimental CAR T cell studies, and is associated with high cytokine release with CAR T cell therapy (henceforth referred to as “aggressive osteosarcoma”)^23,24^. These models were chosen to control for tumor location (DIPG is within the CNS, ALL is primarily outside of but can involve the CNS, and osteosarcoma is exclusively outside of the CNS), CAR T cell target (GD2, CD19 or B7H3), and degree of cytokine release associated with tumor clearance (Figures 1A-1E). Pairs of tumor type and on-target therapeutic (tumor-clearing) CAR T cell are as follows: DIPG with GD2-CAR T^19^, ALL with CD19-CAR T^25^, aggressive osteosarcoma (engineered to express CD19)^24^ with CD19-CAR T, and rapid-clearing osteosarcoma with either GD2- or B7H3-CAR T. All CAR T cell constructs contain the 4-1BBz co-stimulatory domains^24,26^. Tumor clearance timelines varied across models and are as shown in Figures 1B-1E. All experiments were performed with a mock T cell control, for which tumor-bearing mice received T cells which were identically manipulated and activated but do not express a CAR construct. In the DIPG and ALL models, we included an additional off-target CAR T-cell control group that does not treat the tumor in that model and controls for possible effects of these CAR T cells (CD19- and GD2-targeting) on normal tissues.

**Figure 1.**
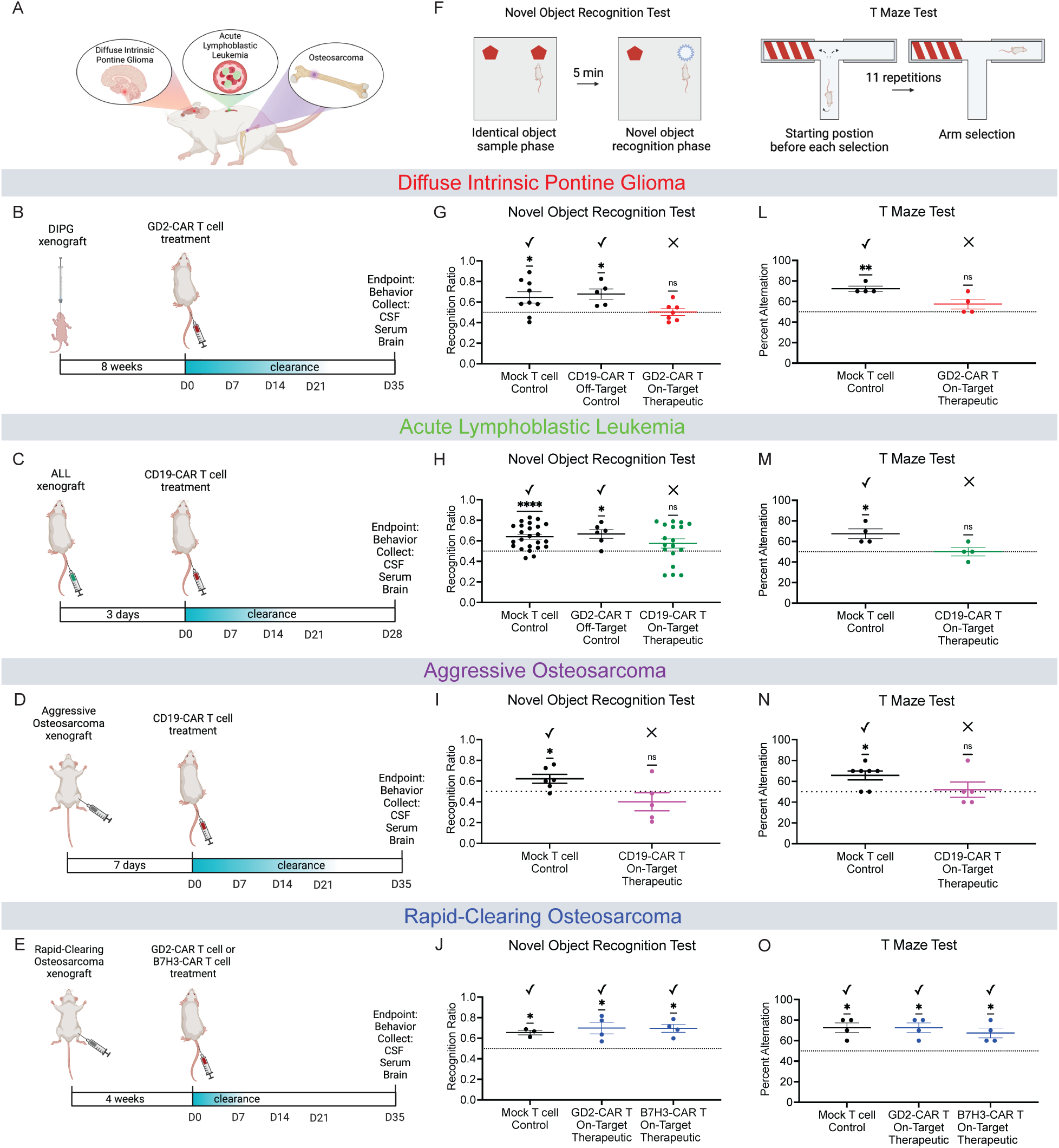
Cognitive deficits and neuroinflammation following tumor-clearing CAR T cell therapy. (A) Schematic illustration denoting the murine xenograft models assessed. (B, C, D, E) Schematic illustration of the various xenograft and CAR T cell therapy experimental timelines. (F) Schematic illustration of the Novel Object Recognition Test (NORT) to evaluate attention and short-term memory. Recognition ratio is defined as the percent time spent with the novel object over the total time spent interacting with either object. (G) DIPG xenografted mice treated with either mock T cells, Off-Target CD19-CAR T cells, or On-Target Therapeutic GD2-CAR T cells. NORT performed on day 34 post-CAR T therapy. Mock T cell Control (n=9 mice), CD19-CAR T Control (n=5 mice), GD2-CAR T Therapeutic (n=7 mice). (H) ALL xenografted mice treated with either mock T cells, Off-Target GD2-CAR T cells, or On-Target Therapeutic CD19-CAR T cells. NORT performed on day 27 post-CAR T therapy. Mock T cell Control (n=24 mice), GD2-CAR T Control (n=6 mice), CD19-CAR T Therapeutic (n=18 mice). (I) Aggressive osteosarcoma xenografted mice treated with either mock T cells or On-Target Therapeutic CD19-CAR T cells. NORT performed on day P34 post-CAR T therapy. Mock T cell Control (n=6 mice), CD19-CAR T Therapeutic (n=5 mice). (J) Rapid-clearing osteosarcoma xenografted mice treated with either mock T cells, On-Target Therapeutic GD2-CAR T, or On-Target Therapeutic B7H3-CAR T cells. NORT performed on day 34 post-CAR T therapy. Mock T cell Control (n=3 mice), GD2-CAR T Therapeutic (n=4 mice), B7H3-CAR T Therapeutic (n=4 mice). (K) T Maze Alternation Test to assess spatial working memory and hippocampal-related cognitive dysfunction. Percent Alternation is calculated over 11 trials allowing for 10 decisions. (L) DIPG xenografted mice treated with either mock T cell Control or On-Target Therapeutic GD2-CAR T cells. T maze performed on day 35 post-CAR T therapy. Mock T cell control (n=4 mice), GD2-CAR T therapeutic (n=4 mice). (M) ALL xenografted mice treated with either mock T cell Control or On-Target Therapeutic CD19-CAR T cells. T maze performed on day 28 post-CAR T therapy. Mock T cell control (n=4 mice), CD19-CAR T therapeutic (n=4 mice). (N) Aggressive osteosarcoma xenografted mice treated with either mock T cell Control or On-Target Therapeutic CD19-CAR T cells. T maze performed on day 35 post-CAR T therapy. Mock T cell control (n=7 mice), CD19-CAR T therapeutic (n=5 mice). (O) Rapid-clearing osteosarcoma xenografted mice treated with either mock T cells, On-Target Therapeutic GD2-CAR T, or On-Target Therapeutic B7H3-CAR T cells. T Maze performed on day 35 post-CAR T therapy. Mock T cell Control (n=4 mice), GD2-CAR T Therapeutic (n=4 mice), B7H3-CAR T Therapeutic (n=4 mice). (G-J and L-O) Data shown as mean ± SEM. Each point = one mouse. ns = p > 0.05, *p < 0.05, **p < 0.01, ****p < 0.0001. Analyzed via One Sample T and Wilcoxon Test.

Early clinical reports suggest that a substantial fraction of patients exhibit long-term neurological sequelae by ∼one year after CAR T cell therapy, including difficulties with attention and memory^3,4^. We therefore tested these cognitive functions in our murine models using a modified novel object recognition test (NORT) to assess for deficits in attention and short term (<5min) memory (Figure 1F). Mice treated with tumor-clearing CAR T cells in the patient-derived DIPG model, ALL model, and aggressive osteosarcoma model each exhibited impaired behavioral performance in the novel object recognition test, indicating attention and/or memory deficits (Figure 1G-1I). In contrast, tumor-bearing mice treated with off-target GD2-CAR T cells, off-target CD19-CAR T cells, or mock T cells, exhibit intact behavioral performance in novel object recognition testing (Figure 1G-1I).

Notably, in the second osteosarcoma model that clears rapidly with CAR T cell therapy, mice treated with either tumor-clearing GD2- or B7H3-CAR T cells demonstrate no cognitive behavioral deficits in novel object recognition testing (Figure 1J), suggesting intact cognition following this relatively efficient CAR T cell-mediated tumor clearance.

To further investigate cognitive performance and specifically assess hippocampal-related spatial working memory, we used the Spontaneous Alternation T Maze Test (Figure 1K). This task is a well-established test of spatial memory that requires intact hippocampal function^27,28^. We found that mice bearing xenografts of DIPG, ALL, or aggressive osteosarcoma and treated with tumor-clearing CAR T cells were unable to adequately perform the T Maze Test (Figure1L-1N). Tumor-bearing mice treated with mock T cells did exhibit adequate performance in the T Maze test (Figures 1L-1O), suggesting that the presence of tumor alone does not account for impaired hippocampal-dependent spatial memory function. In contrast to the cognitive performance in the first three models, the rapid-clearing osteosarcoma model shows intact cognitive performance in the T Maze after tumor-clearing CAR T cell therapy (Figure 1O), consistent with the NORT results above and underscoring that cognitive impairment is not associated with tumor-clearing CAR T cell therapy in every case. Taken together, these NORT and T Maze behavioral data demonstrate impaired cognitive function in both CNS and non-CNS cancer models following tumor-clearing CAR T cell therapy (Figures 1G-1I and 1L-1N) that is not explained by the presence of tumor nor on-target, off-tumor effects of the CAR T cells.

### Persistent CSF cytokine and chemokine elevation after tumor-clearing CAR T cell therapy

To assess the blood and CSF cytokine and chemokine profiles in each tumor model following CAR T cell therapy, we collected cerebrospinal fluid (CSF) ∼one month following therapeutic CAR T cell or mock T cell control administration and measured cytokines and chemokines. At one month following tumor-clearing CAR T cell administration, CSF cytokines were elevated above the levels seen in tumor-bearing, mock T cell-treated controls in the DIPG, ALL and aggressive osteosarcoma models (Figure 2A-2C), but not in the rapid-clearing osteosarcoma model (Figure 2D). Prominently, we observed persistently elevated levels of eotaxin/CCL11, CCL2, CCL7, CXCL1, CXCL10 and BAFF in the DIPG, ALL, and aggressive osteosarcoma models (Figure 2A-2C), the same three models that also exhibit cognitive deficits (Figure 1). Eotaxin/CCL11 is of particular interest as it is a chemokine sufficient to cause cognitive impairment, microglial reactivity in the hippocampus and reduced hippocampal neurogenesis^15,29^. To better understand why mice treated for rapid-clearing osteosarcoma may not exhibit elevated cytokine/chemokine levels in the CSF we also examined serum cytokines and chemokines in the two osteosarcoma models (Fig S1). At one month following tumor-clearing CAR T cell therapy, the aggressive osteosarcoma model exhibited persistently elevated serum levels of CCL11, CCL2 and BAFF compared to tumor-bearing, mock T cell-treated control mice (Fig S1A-S1B). In contrast, the rapid-clearing osteosarcoma model does not exhibit significantly elevated levels of any measured cytokine (Fig S1A-S1B) in the serum. This raises the possibility that CAR T cell therapy in this cognitively intact model does not elicit a similar immune response during the CAR T cell-mediated tumor-clearing process as the cognitively impaired models, suggesting that it is not the CAR T cell-tumor interaction per se but rather a broader accompanying immune response that triggers CNS immune dysregulation.

**Figure 2.**
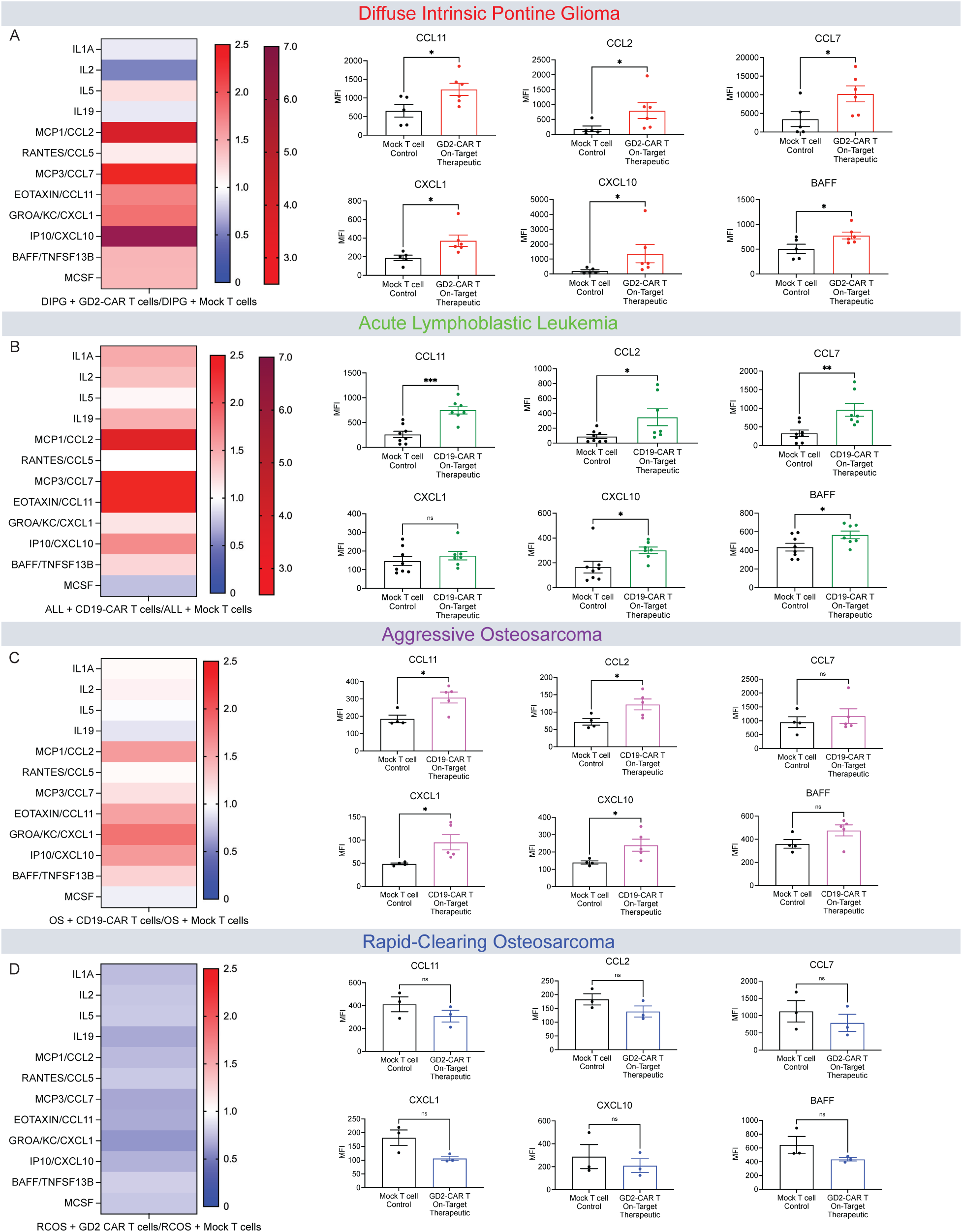
Persistent CSF cytokine and chemokine elevation after tumor-clearing CAR T cell therapy. (A) Left, heat map of fold change of cytokine and chemokine analysis of CSF in DIPG+On-Target Therapeutic GD2-CAR T cell treated mice relative to DIPG+mock T cell treated control mice 35-days post-CAR T cell therapy. Right, quantification of raw MFI of CSF levels of cytokines (CCL11, CCL2, CCL7, CXCL1, CXCL10, BAFF). (B) Left, heat map of fold change of cytokine and chemokine analysis of CSF in ALL+On-Target Therapeutic CD19-CAR T cell treated mice relative to ALL+mock T cell treated control mice 28-days post-CAR T cell therapy. Right, quantification of raw MFI of CSF levels of cytokines (CCL11, CCL2, CCL7, CXCL1, CXCL10, BAFF). (C) Left, heat map of fold change of cytokine and chemokine analysis of CSF in aggressive osteosarcoma (OS)+On-Target Therapeutic CD19-CAR T cell treated mice relative to aggressive osteosarcoma (OS)+mock T cell treated control mice 35-days post-CAR T cell therapy. Right, quantification of raw MFI of CSF levels of cytokines (CCL11, CCL2, CCL7, CXCL1, CXCL10, BAFF). (D) Left, heat map of fold change of cytokine and chemokine analysis of CSF in rapid-clearing osteosarcoma (RCOS)+On-Target Therapeutic GD2-CAR T cell treated mice relative to rapid-clearing osteosarcoma (RCOS)+mock T cell treated control mice 35-days post-CAR T cell therapy. Right, quantification of raw MFI of CSF levels of cytokines (CCL11, CCL2, CCL7, CXCL1, CXCL10, BAFF). (A-D) Cytokine and chemokines significantly elevated or depressed, analyzed via Student’s T test of raw MFI values.

### Persistent white matter microglial reactivity after tumor-clearing CAR T cell therapy

Previous work has demonstrated that a wide variety of both CNS-directed and systemic insults can induce reactivity of microglia, the CNS-resident tissue macrophages, especially in white matter regions^8,11,15^. We therefore set out to evaluate the presence and distribution of microglial reactivity following CAR T cell therapy in the various CAR T cell therapy models used here. We assessed microglial number (assessed by positivity for the pan-microglial marker Iba1) and reactivity (assessed by Iba1, CD68 co-positivity) of microglia in the cortex, subcortical white matter, and white matter of the hippocampal dentate gyrus. Both microglial numbers and reactivity were increased in subcortical white matter and hippocampal white matter after tumor-clearing CAR T cell therapy for DIPG xenografted to the brainstem (Figures 3A-3B; S2A-S2B). Microglial number but not reactivity was also increased in the cortex after tumor-clearing CAR T cell therapy for DIPG (Figure 3C; S2C). Following tumor-clearing CAR T cell therapy for ALL, we found increased microglial reactivity in the subcortical white matter, increased microglial numbers and reactivity in the hippocampal white matter, and no changes in cortical microglia (Figures 3D-3F; S2D-S2F). The aggressive osteosarcoma model exhibits an increase in microglial number and reactivity in subcortical white matter, increased microglial reactivity in the hippocampal white matter, and no changes in cortical microglia following tumor-clearing CAR T cell therapy (Figures 3G-3I; S2G-S2I). In the rapid-clearing osteosarcoma model with intact cognitive behavioral performance, neither GD2-nor B7H3-targeting, tumor-clearing CAR T cell therapy resulted in an increase in microglial number or reactivity in any of the regions evaluated (Figure 3J-3L; S2J-S2L). Microglia were not elevated in off-target control groups in any model, indicating that on-target, off-tumor effects of the CAR T cells do not account for the microglial changes observed in the tumor-clearing CAR T cell groups. Taken together, these data illustrate persistent reactivity chiefly involving white matter microglia following tumor-clearing CAR T cell therapy in the mouse models that also exhibit cognitive behavioral deficits and CSF cytokine/chemokine elevation.

**Figure 3.**
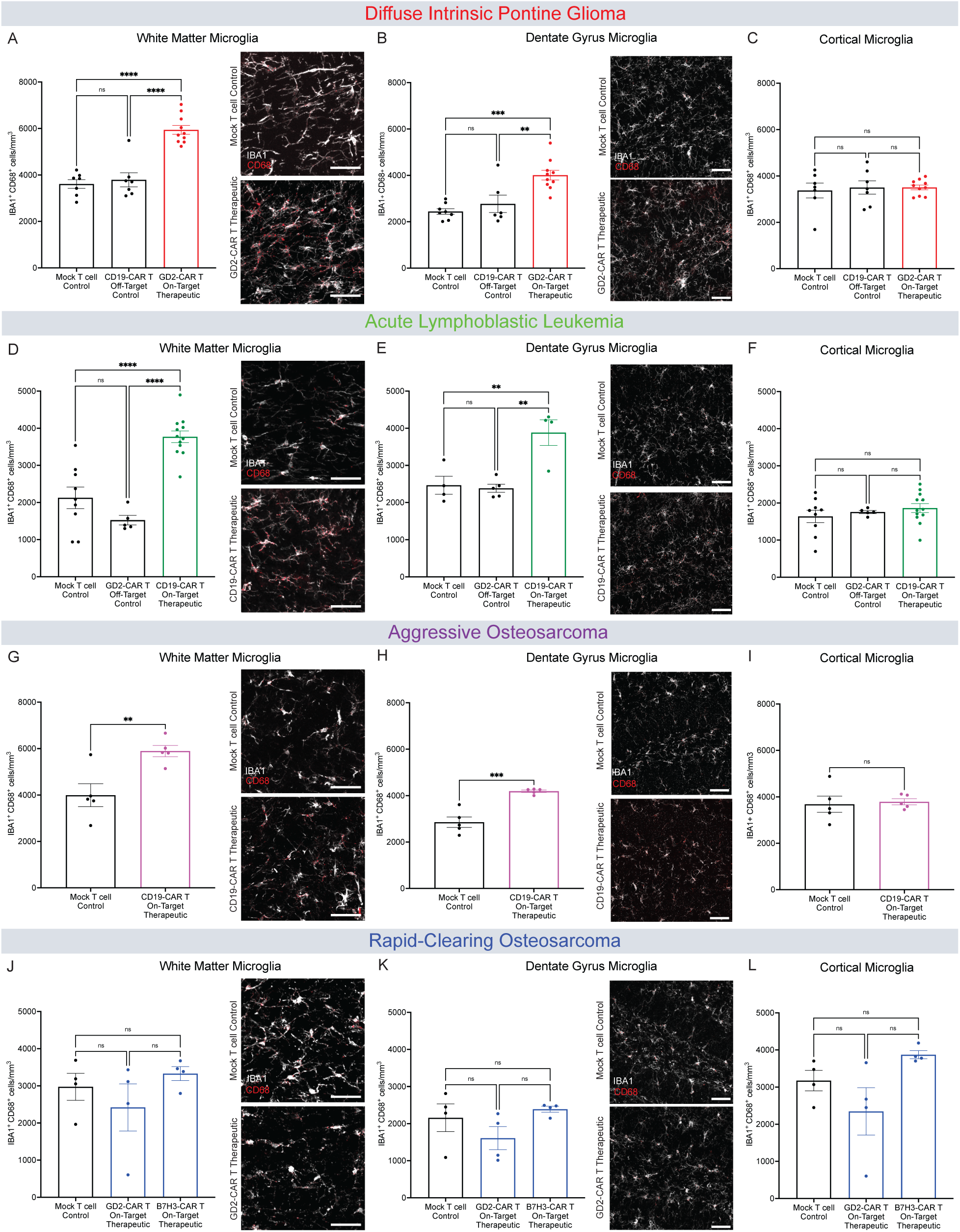
Tumor-clearing CAR T Cell therapy can result in white matter microglial reactivity. (A) Quantification of DIPG xenografted mice microglial reactivity (IBA1+ CD68+) and representative confocal micrographs (white = IBA1; red = CD68) from the corpus callosum 35 days post-CAR T therapy. Mock T cell Control (n=7 mice), CD19-CAR T Control (n=7 mice), GD2-CAR T Therapeutic (n=10 mice). (B) Quantification of DIPG xenografted mice microglial reactivity (IBA1+ CD68+) and representative confocal micrographs (white = IBA1; red = CD68) 35 days post-CAR T cell therapy in the hilar white matter region of the dentate gyrus. Mock T cell Control (n=8 mice), CD19-CAR T Control (n=6 mice), GD2-CAR T Therapeutic (n=10 mice). (C) Quantification of DIPG xenografted mice microglial reactivity (IBA1+ CD68+) 35 days post-CAR T therapy in the cortical grey matter. Mock T cell Control (n=7 mice), CD19-CAR T Control (n=7 mice), GD2-CAR T Therapeutic (n=10 mice). (D) Quantification of ALL xenografted mice microglial reactivity (IBA1+ CD68+) and representative confocal micrographs (white = IBA1; red = CD68) from the corpus callosum 28 days post-CAR T therapy. Mock T cell Control (n=9 mice), GD2-CAR T Control (n=5 mice), CD19-CAR T Therapeutic (n=12 mice). (E) Quantification of ALL xenografted mice microglial reactivity (IBA1+ CD68+) and representative confocal micrographs (white = IBA1; red = CD68;) 28 days post-CAR T cell therapy in the hilar white matter region of the dentate gyrus. Mock T cell Control (n=4 mice), GD2-CAR T Control (n=5 mice), CD19-CAR T Therapeutic (n=4 mice). (F) Quantification of ALL xenografted mice microglial reactivity (IBA1+ CD68+) 28 days post-CAR T therapy in the cortical grey matter. Mock T cell Control (n=9 mice), GD2-CAR T Control (n=5 mice), CD19-CAR T Therapeutic (n=12 mice). (G) Quantification of aggressive osteosarcoma xenografted mice microglial reactivity (IBA1+ CD68+) and representative confocal micrographs (white = IBA1; red = CD68) from the corpus callosum 35 days post-CAR T therapy. Mock T cell Control (n=5 mice), CD19-CAR T Therapeutic (n=5 mice). (H) Quantification of aggressive osteosarcoma xenografted mice microglial reactivity (IBA1+ CD68+) and representative confocal micrographs (white = IBA1; red = CD68) 35 days post-CAR T cell therapy in the hilar white matter region of the dentate gyrus. Mock T cell Control (n=5 mice), CD19-CAR T Therapeutic (n=5 mice). (I) Quantification of aggressive osteosarcoma xenografted mice microglial reactivity (IBA1+ CD68+) 35 days post-CAR T therapy in the cortical grey matter. Mock T cell Control (n=5 mice), CD19-CAR T Therapeutic (n=5 mice). (J) Quantification of rapid-clearing osteosarcoma xenografted mice microglial reactivity (IBA1+ CD68+) and representative confocal micrographs (white = IBA1; red = CD68) from the corpus callosum 35 days post-CAR T therapy. Mock T cell Control (n=4 mice), GD2-CAR Therapeutic (n=4 mice), B7H3-CAR T Therapeutic (n=4 mice). (K) Quantification of rapid-clearing osteosarcoma xenografted mice microglial reactivity (IBA1+ CD68+) and representative confocal micrographs (white = IBA1; magenta = CD68) 35 days post-CAR T cell therapy in the hilar white matter region of the dentate gyrus. Mock T cell Control (n=4 mice), GD2-CAR T Therapeutic (n=4 mice), B7H3-CAR T Therapeutic (n=4 mice). (L) Quantification of rapid-clearing osteosarcoma xenografted mice microglial reactivity (IBA1+ CD68+) 35 days post-CAR T therapy in the cortical grey matter. Mock T cell Control (n=4 mice), GD2-CAR Therapeutic (n=4 mice), B7H3-CAR T Therapeutic (n=4 mice). (A-F, J-L) Data shown as mean ± SEM. Each point = one mouse, ns = p > 0.05, **p < 0.01, ***p < 0.001, ****p < 0.0001, analyzed via One-way ANOVA. (G-I) Data shown as mean ± SEM. Each point = one mouse, ns = p > 0.05, **p < 0.01, analyzed via Unpaired T Test. Scale bars equal 50um in all confocal images.

### Tumor-clearing CAR T cell therapy can disrupt oligodendroglial lineage cell dynamics

Microglial reactivity can dysregulate oligodendroglial homeostasis and plasticity^11,12^. To assess oligodendroglial integrity after CAR T cell therapy, we quantified oligodendrocyte precursor cells (OPCs) and mature oligodendrocytes in the cortex and subcortical white matter. In the cortex, we found no changes after tumor-clearing or off-target CAR T cell administration in any of the models (Figure S3A-S3H). However, subcortical white matter exhibited a marked decrease in both OPCs (20-25% decrease, Figures 4A-4C) and oligodendrocytes (25-35% decrease, Figures 4D-4F) at one month following tumor-clearing CAR T cell administration in each of the DIPG, ALL and aggressive osteosarcoma models, compared to tumor-bearing mock T cell and off-target CAR T cell controls (Figures 4A-4F). The rapid-clearing osteosarcoma model showed no changes in OPCs or oligodendrocytes after tumor-clearing GD2 or B7H3-CAR T cell treatment (Figures 4G and 4H), concordant with the lack of microglial reactivity, CSF cytokine elevation or cognitive behavioral deficits in this model as described above.

**Figure 4.**
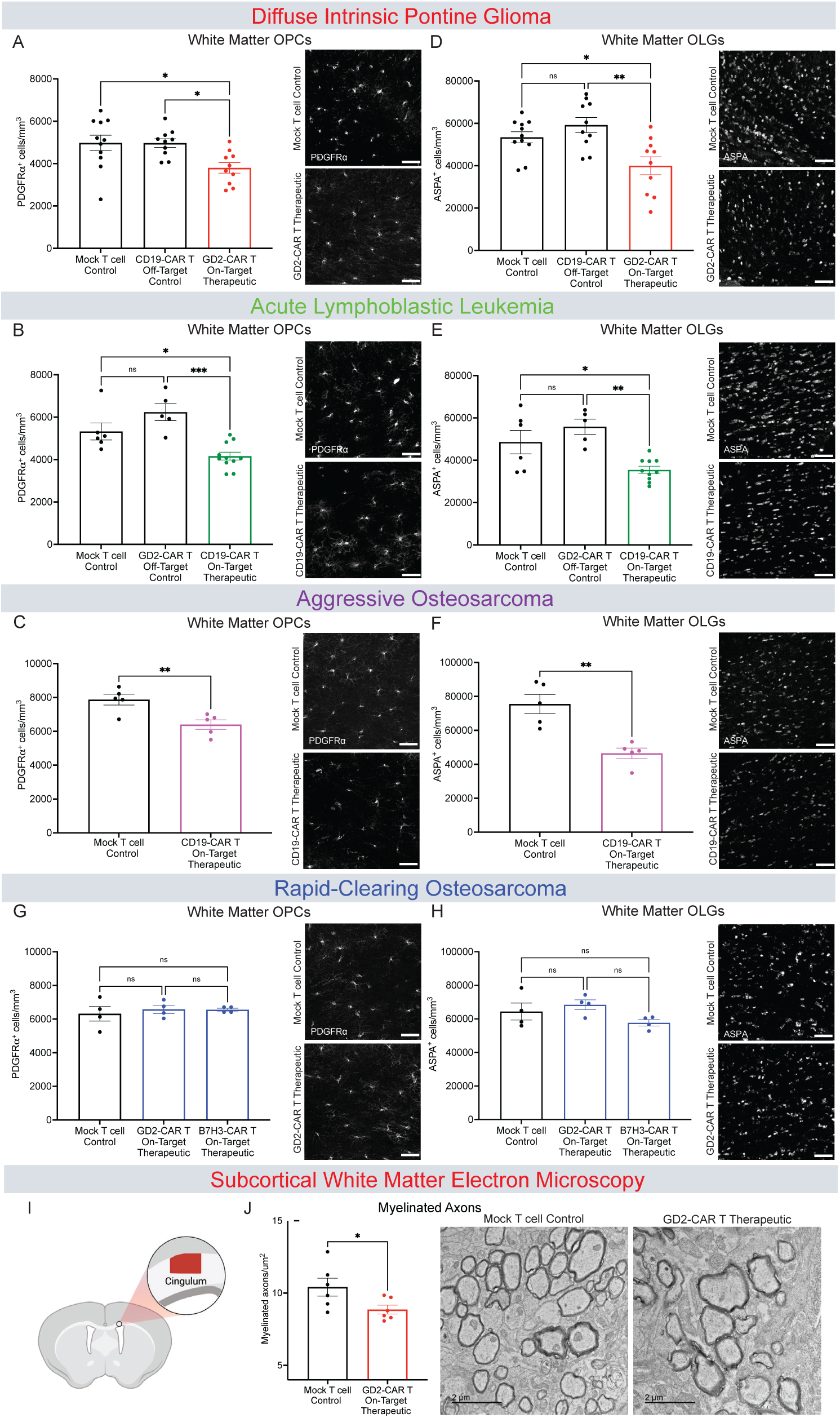
Tumor-clearing CAR T Cell therapy can result in oligodendroglial and myelin deficits. (A) Quantification and representative confocal micrographs of PDGFRa+ oligodendrocyte precursor cells from the corpus callosum in DIPG xenografted mice 35 days post-CAR T therapy. Mock T cell Control (n=11 mice), CD19-CAR T Control (n=10 mice), GD2-CAR T Therapeutic (n=10 mice). (B) Quantification and representative confocal micrographs of PDGFRa+ oligodendrocyte precursor cells from the corpus callosum in ALL xenografted mice 28 days post-CAR T therapy. Mock T cell Control (n=6 mice), GD2-CAR T Control (n=5 mice), CD19-CAR T Therapeutic (n=11 mice). (C) Quantification and representative confocal micrographs of PDGFRa+ oligodendrocyte precursor cells from the corpus callosum in aggressive osteosarcoma xenografted mice 35 days post-CAR T therapy. Mock T cell Control (n=5 mice), CD19-CAR T Therapeutic (n=5 mice). (D) Quantification and representative confocal micrographs of ASPA+ mature oligodendrocytes from corpus callosum in DIPG xenografted mice 35 days post-CAR T therapy. Mock T cell Control (n=11 mice), CD19-CAR T Control (n=10 mice), GD2-CAR T Therapeutic (n=10 mice). (E) Quantification and representative confocal micrographs of ASPA+ mature oligodendrocytes from the corpus callosum in ALL xenografted mice 28 days post-CAR T therapy. Mock T cell Control (n=6 mice), CD19-CAR T Control (n=5 mice), GD2-CAR T Therapeutic (n=10 mice). (F) Quantification and representative confocal micrographs of ASPA+ mature oligodendrocytes from the corpus callosum in aggressive osteosarcoma xenografted mice 35 days post-CAR T therapy. Mock T cell Control (n=5 mice), CD19-CAR T Therapeutic (n=5 mice). (G) Quantification and representative confocal micrographs of PDGFRa+ oligodendrocyte precursor cells from the corpus callosum in rapid-clearing osteosarcoma xenografted mice 35 days post-CAR T therapy. Mock T cell Control (n=4 mice), GD2-CAR T Therapeutic l (n=4 mice), B7H3-CAR T Therapeutic (n=4 mice). (H) Quantification and representative confocal micrographs of ASPA+ mature oligodendrocytes from the corpus callosum in rapid-clearing osteosarcoma xenografted mice 35 days post-CAR T therapy. Mock T cell Control (n=4 mice), GD2-CAR T Therapeutic (n=4 mice), B7H3-CAR T Therapeutic (n=4 mice). (I) Schematic illustration denoting the cingulum region of the corpus callosum used for the transmission electron microscopy (TEM) DIPG xenografted mice 35 days post-CAR T therapy. (J) Quantification and Representative TEM images of white matter myelinated axons per area at day P35 post-CAR T therapy. Myelinated axonal projections are in cross section and visible as electron-dense sheaths encircling axons. Scale bar = 2 um. Mock T cell control (n=6 mice), GD2-CAR T therapeutic (n=6 mice). (A, B, D, E, G and H) Data shown as mean ± SEM. Each point = one mouse. ns = p > 0.05, *p < 0.05, **p < 0.01, ***p < 0.001, analyzed via One-way ANOVA. (C, F and J) Data shown as mean ± SEM. Each point = one mouse, *p < 0.05, analyzed via Unpaired T Test. Scale bars equal 50um in all confocal micrographs (A-H), and 2um in electron microscopy images (J).

This reduction in mature oligodendrocytes suggests potential disruption of myelin homeostasis. To assess myelin ultrastructure, we performed transmission electron microscopy (TEM) of the subcortical white matter (cingulum of the corpus callosum). We found a ∼20% decrease in subcortical white matter myelinated axon density following GD2-CAR T cell clearance of DIPG xenografted to the brainstem in comparison to mock T cell-treated, tumor-bearing controls (Figures 4I-4J). No significant difference was observed in myelin sheath thickness (*g*-ratio) between groups (Figure S3I-S3J).

### Tumor-clearing CAR T cell therapy can impair hippocampal neurogenesis

Reactive microglia and particular cytokines/chemokines, including CCL11, can impair hippocampal neurogenesis^8,15,29,30^, an important component of neural plasticity that contributes to spatial memory function^31–34^. Given the observed increase in dentate gyrus white matter reactive microglia and in CCL11 and other CSF cytokines/chemokines, together with spatial memory deficits, we hypothesized that neurogenesis may be impaired following tumor-clearing CAR T cell administration.

We next investigated neurogenesis in the dentate gyrus of the hippocampus by quantifying doublecortin-positive neuroblasts^35,36^. We found that neurogenesis is diminished by 40-50% following tumor-clearing CAR T cell therapy in the DIPG, ALL, and aggressive osteosarcoma models, compared to off-target CAR T or mock T cell controls (Figure 5A-5C). Interestingly, the rapid-clearing osteosarcoma model exhibited a small increase in neurogenesis following tumor-clearing B7H3-CAR T cell administration, but was at control levels following tumor-clearing GD2-CAR T cell administration (Figure 5D). Taken together, these findings are concordant with the emerging pattern of multicellular dysregulation and cognitive impairment following CAR T cell therapy that induces persistent neuroinflammation.

**Figure 5.**
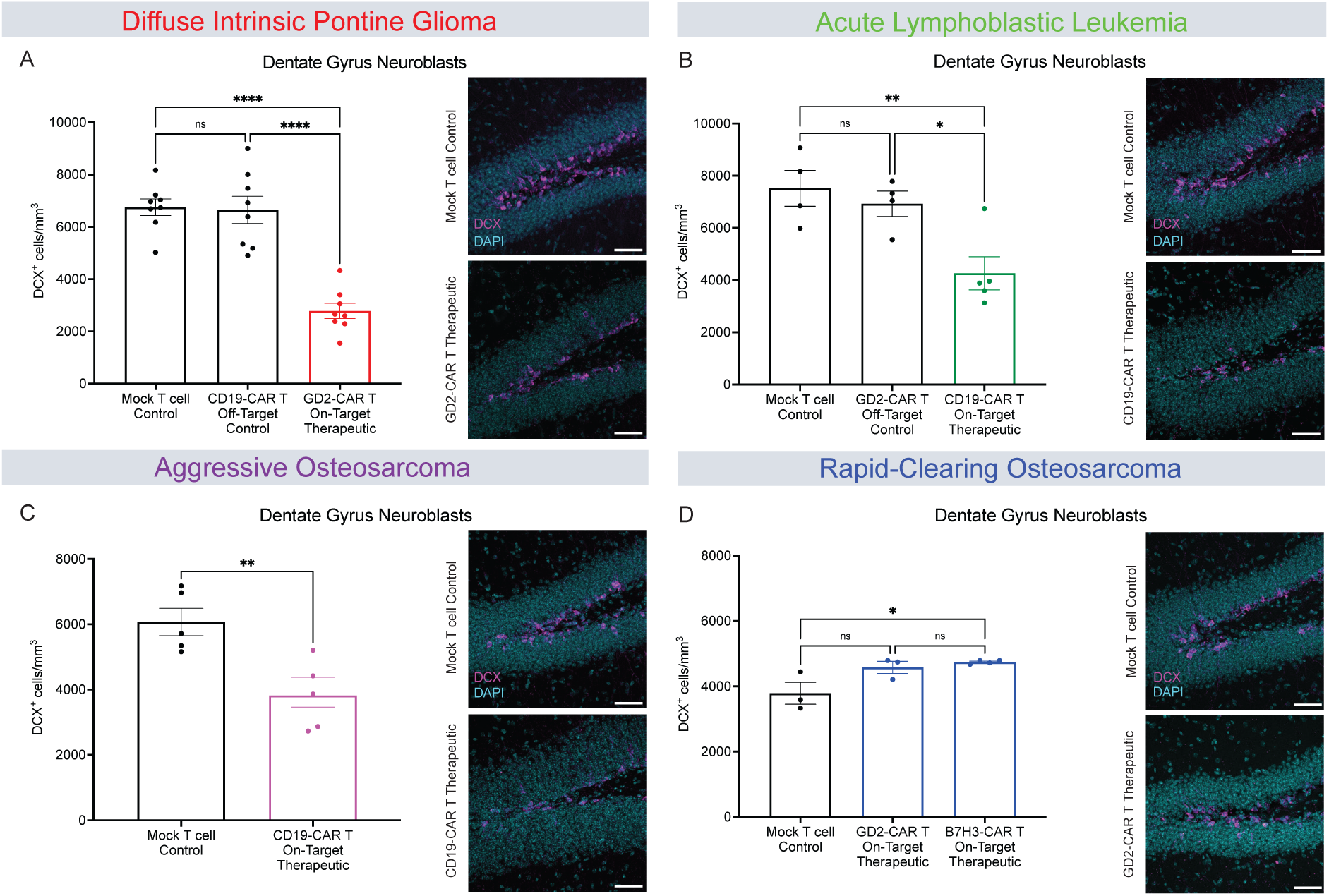
Tumor-clearing CAR T Cell therapy can impair hippocampal neurogenesis. (A) Quantification of neuroblasts (DCX+) in DIPG model and representative confocal micrographs (magenta = DCX; cyan = DAPI) from the dentate gyrus 35 days after CAR T cell therapy. Mock T cell control (n=8 mice), CD19-CAR T control (n=8 mice), GD2-CAR T Therapeutic (n=8 mice). (B) Quantification of neuroblasts (DCX+) in ALL model and representative confocal micrographs (magenta = DCX; cyan = DAPI) from the dentate gyrus 28 days after CAR T cell therapy. Mock T cell control (n=4 mice), GD2-CAR T control (n=4 mice), CD19-CAR T Therapeutic (n=5 mice). (C) Quantification of neuroblasts (DCX+) in aggressive osteosarcoma model and representative confocal micrographs (magenta = DCX; cyan = DAPI) from the dentate gyrus 35 days after CAR T cell therapy. Mock T cell control (n=5 mice), CD19-CAR T Therapeutic (n=5 mice). (D) Quantification of neuroblasts (DCX+) in rapid-clearing osteosarcoma model and representative confocal micrographs (magenta = DCX; cyan = DAPI) from the dentate gyrus 35 days after CAR T cell therapy. Mock T cell control (n=3 mice), GD2-CAR T Therapeutic (n=3 mice), B7H3-CAR T Therapeutic (n=4 mice). (A, B, and D) Data shown as mean ± SEM. Each point = one mouse. ns = p > 0.05, *p < 0.05, **p < 0.01, ****p < 0.0001, analyzed via One-way ANOVA. (C) Data shown as mean ± SEM. Each point = one mouse, **p < 0.01, analyzed via Unpaired T Test. Scale bars equal 50um in all confocal images.

### Anti-cytokine therapy targeting IL1a does not rescue cognitive performance

Anti-cytokine agents such as Anakinra, an interleukin-1 receptor antagonist, are frequently administered to CAR T cell patients to treat acute neurotoxicity such as immune effector cell-associated neurotoxicity (ICANS) frequently seen in patients with ALL^37,38^. We therefore assessed the possible effects of Anakinra on the observed cellular dysregulation and cognitive deficits in the ALL model to assess possible rescue of cellular homeostasis and cognitive behavior after tumor-clearing CAR T cell therapy. Anakinra was administered (10 mg/kg i.p.) from day 0 to day 14 of CAR T cell treatment (Figure S4A). We found no effects on Anakinra on microglial reactivity, oligodendroglial deficits, hippocampal neurogenesis deficits nor rescue of cognitive performance (Figures S4B-S4F).

### CSF1R inhibition differentially depletes microglia in subcortical and hippocampal white matter

Microglial reactivity and consequent dysregulation of oligodendroglial lineage cells and hippocampal neurogenesis is thought to be central to cognitive impairment following traditional cancer therapies such as radiation and chemotherapy (for review see^17^). Accordingly, microglial depletion rescues cellular and cognitive deficits after traditional cancer therapies^11,39^. Therefore, we tested whether microglial depletion after the period of CAR T cell-mediated tumor clearance could rescue cellular and cognitive impairment in these mouse models.

To deplete microglia, we used a small-molecule inhibitor of Colony Stimulating Factor 1 Receptor (CSF1R), PLX5622^40^. Most microglia and macrophages depend on CSF1R signaling for survival and therefore, treatment with the CSF1R inhibitor (CSF1Ri) results in swift clearance of the majority of microglia in the brain^41^. Not all microglia are CSF1R-dependent, and a fraction of microglia remain^42^. We administered CSF1Ri formulated in mouse chow from D21 to D35 in the DIPG model (Figure 6A) and from D14 to D28 in the ALL model (Figure 6B). Following 14 days of CSF1R inhibitor treatment, microglia were depleted by 70-75% in subcortical white matter (Figure 6C-6D), but only depleted by 45-55% in the hippocampal white matter (Figures 6E-6F), underscoring the regional heterogeneity of microglial subpopulations.

**Figure 6.**
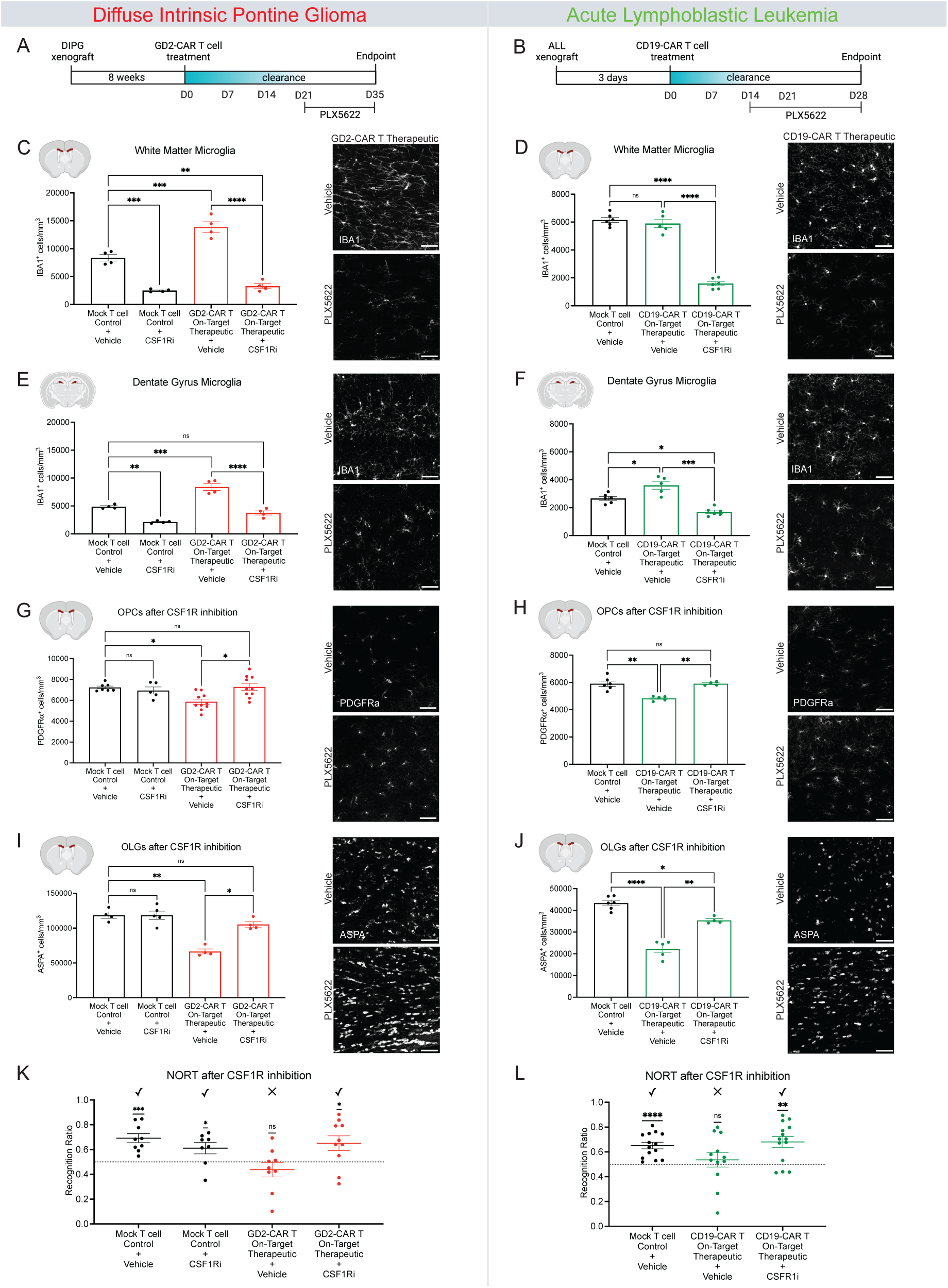
Microglial depletion rescues oligodendroglial dysregulation cognitive performance in novel object recognition testing. (A) Schematic illustrations of experimental timeline of the DIPG model including administration of the CSF1R inhibitor (PLX5622). (B) Schematic illustrations of experimental timeline of the ALL model including administration of the CSF1R inhibitor (PLX5622). (C) Quantification of DIPG xenografted mice corpus callosum white matter microglia (IBA1+) 35 days post-CAR T therapy with or without 14 days of CSF1R inhibition. Mock T cell+Vehicle Chow (n=4 mice), mock T cell+CSF1Ri Chow (n=4 mice), GD2-CAR T Therapeutic+Vehicle Chow (n=4 mice), GD2-CAR T Therapeutic+CSF1Ri Chow (n=4 mice). Representative confocal micrographs of microglia (white = IBA1) in the corpus callosum of DIPG xenografted mice 35 days after CAR T cell therapy with or without 14 days of CSF1R inhibitor (PLX5622). (D) Quantification of ALL xenografted mice corpus callosum white matter microglia (IBA1+) 28 days post-CAR T therapy with or without 14 days of CSF1R inhibition. Mock T cell+Vehicle Chow (n=6 mice), CD19-CAR T Therapeutic+Vehicle Chow (n=5 mice), CD19-CAR T Therapeutic+CSF1Ri Chow (n=6 mice). Representative confocal micrographs of microglia (white = IBA1) in the corpus callosum of ALL xenografted mice 28 days after CAR T cell therapy with or without 14 days of CSF1R inhibitor (PLX5622). (E) Quantification of DIPG xenografted mice dentate gyrus microglia (IBA1+) 35 days post-CAR T therapy with or without 14 days of CSF1R inhibition. Mock T cell+Vehicle Chow (n=4 mice), mock T cell+CSF1Ri Chow (n=4 mice), GD2-CAR T Therapeutic+Vehicle Chow (n=4 mice), GD2-CAR T Therapeutic+CSF1Ri Chow (n=4 mice). Representative confocal micrographs of microglia (white = IBA1) in the dentate gyrus of DIPG xenografted mice 35 days after CAR T cell therapy with or without 14 days of CSF1R inhibitor (PLX5622). (F) Quantification of ALL xenografted mice dentate gyrus microglia (IBA1+) 28 days post-CAR T therapy with or without 14 days of CSF1R inhibition. Mock T cell+Vehicle Chow (n=6 mice), CD19-CAR T Therapeutic+Vehicle Chow (n=5), CD19-CAR T Therapeutic+CSF1Ri Chow (n=6 mice). Representative confocal micrographs of microglia (white = IBA1) in the dentate gyrus of ALL xenografted mice 28 days after CAR T cell therapy with or without 14 days of CSF1R inhibitor (PLX5622). (G) Quantification of DIPG xenografted mice oligodendrocyte precursor cells (PDGFRa+) 35 days post-CAR T therapy with or without 14 days of CSF1R inhibition. Mock T cell+Vehicle Chow (n=7 mice), mock T cell+CSF1Ri Chow (n=5 mice), GD2-CAR T Therapeutic+Vehicle Chow (n=10 mice), GD2-CAR T Therapeutic+CSF1Ri Chow (n=10 mice). Representative confocal micrographs of PDGFRa+ oligodendrocyte precursor cells (white = PDGFRa) in the corpus callosum of DIPG xenografted mice 35 days after CAR T cell therapy with or without 14 days of CSF1R inhibitor (PLX5622). (H) Quantification of ALL xenografted mice oligodendrocyte precursor cells (PDGFRa+) 28 days post-CAR T therapy with or without 14 days of CSF1R inhibition. Mock T cell+Vehicle Chow (n=6 mice), CD19-CAR T Therapeutic+Vehicle Chow (n=5 mice), CD19-CAR T Therapeutic+CSF1Ri Chow (n=4 mice). Representative confocal micrographs of PDGFRa+ oligodendrocyte precursor cells (white = PDGFRa) in the corpus callosum of ALL xenografted mice 28 days after CAR T cell therapy with or without 14 days of CSF1R inhibitor (PLX5622). (I) Quantification of DIPG xenografted mice mature oligodendrocytes (ASPA+) 35 days post-CAR T therapy with or without 14 days of CSF1R inhibition. Mock T cell+Vehicle Chow (n=4 mice), mock T cell+CSF1Ri Chow (n=5 mice), GD2-CAR T Therapeutic+Vehicle Chow (n=4 mice), GD2-CAR T Therapeutic+CSF1Ri Chow (n=4 mice). Representative confocal micrographs of ASPA+ oligodendrocytes (white = ASPA) in the corpus callosum of DIPG xenografted mice 35 days after CAR T cell therapy with or without 14 days of CSF1R inhibitor (PLX5622). (J) Quantification of ALL xenografted mice mature oligodendrocytes (ASPA+) 28 days post-CAR T therapy with or without 14 days of CSF1R inhibition. Mock T cell+Vehicle Chow (n=6 mice), CD19-CAR T Therapeutic+Vehicle Chow (n=5 mice), CD19-CAR T Therapeutic+CSF1Ri Chow (n=4 mice). Representative confocal micrographs of ASPA+ oligodendrocytes (white = ASPA) in the corpus callosum of ALL xenografted mice 28 days after CAR T cell therapy with or without 14 days of CSF1R inhibitor (PLX5622). (K) NORT results from DIPG xenografted mice 34 days post-CAR T therapy with or without 14 days of CSF1R inhibition. Mock T cell+Vehicle Chow (n=9 mice), mock T cell+CSF1Ri Chow (n=8 mice), GD2-CAR T Therapeutic+Vehicle Chow (n=9 mice), GD2-CAR T Therapeutic+CSF1Ri Chow (n=10 mice). (L) NORT results from ALL xenografted mice 27 days post-CAR T therapy with or without 14 days of CSF1R inhibition. Mock T cell+Vehicle Chow (n=14 mice), CD19-CAR T Therapeutic+Vehicle Chow (n=12 mice), CD19-CAR T Therapeutic+CSF1Ri Chow (n=14 mice). Data shown as mean ± SEM (C-J). Each point = one mouse. ns = p > 0.05, * p < 0.05, **p < 0.01, ***p < 0.001, ****p < 0.0001, analyzed via 2-way ANOVA. (K and L) Data shown as mean ± SEM. Each point = one mouse. ns = p > 0.05, *p < 0.05, **p < 0.01, ***p < 0.001, ****p < 0.0001. Analyzed via One Sample T and Wilcoxon Test. Scale bars equal 50um in all confocal images (A-J).

In the context of this partial microglial depletion, neither hippocampal neurogenesis nor spatial memory performance were rescued by CSF1R inhibition following tumor-clearing CAR T cell therapy for DIPG or ALL (Figure S5).

### Microglial depletion rescues oligodendroglial dysregulation and cognitive behavior

Given the more robust depletion of subcortical microglia with CSF1R inhibition, we next tested the role of reactive microglia in the subcortical oligodendroglial dysfunction observed following tumor-clearing CAR T cell therapy. We found that subcortical microglial depletion led to a normalization of oligodendroglial lineage cell populations, with a rescue of OPC and mature oligodendrocyte numbers following tumor-clearing CAR T cell administration in both the DIPG and ALL models (Figure 6G-6J). Together with normalization of myelin-forming cells, microglial depletion rescued cognitive performance in the novel object recognition test of attention and short-term memory following tumor-clearing CAR T cell therapy in both DIPG and ALL models (Figures 6K and 6L). In contrast, and consistent with the results above, mice treated with tumor-clearing CAR T cell therapy for either DIPG or ALL who received vehicle did not discern between the novel and familiar object (Figures 6K and 6L). These findings implicate reactive microglia as central to the subcortical oligodendroglial lineage dysregulation and associated cognitive behavioral impairment observed following tumor-clearing CAR T cell therapy.

## Discussion

Taken together, the findings presented here illustrate persistent neuroinflammatory changes, multi-lineage neural cellular dysregulation and consequent cognitive impairment in multiple mouse models of CAR T cell therapy for CNS and non-CNS malignancies. Concordant with early reports of cognitive deficits in patients years following immunotherapy^3–5^ that are clinically similar to those experienced by cancer survivors treated with traditional therapies (for review, see^6^), we find that immunotherapy-related cognitive impairment (IRCI) shares mechanistic underpinnings with cognitive impairment caused by chemotherapy^11–13^ and cranial radiation^7–9^. Both traditional and immune-based therapies elicit a consistent pattern of white matter-selective microglial reactivity, consequent disruption of oligodendroglial homeostasis and hippocampal neurogenesis, and impairment in attention and memory function.

Importantly, control experiments illustrate that on-target, off-tumor effects on normal tissue do not account for the neuroinflammation nor the cellular and behavioral deficits observed in the context of tumor clearance, at least for the specific CAR T cells studied. The GD2 antigen is identical in humans and in mice^43^, thus the non-tumor-targeting GD2-CAR-T cell control groups served to control for on-target, off-tumor effects of the human GD2-CAR T cells used in these mouse models. However, it should be noted that CAR T cell reactivity with antigens on normal neural tissues could occur and cause neurological sequelae with different antigen-targeting CAR T cell therapies, as appears may happen with BCMA-targeting CAR T cells^44^. Additionally, while the presence of a tumor alone can cause a degree of CSF cytokine/chemokine elevation, especially for cancers that involve the CNS, the presence of the tumor alone did not account for the microglial reactivity or downstream cognitive effects observed in any of the models studied.

We find here that while CAR T cell therapy can cause neuroinflammation that triggers a cascade of downstream effects resulting in cognitive dysfunction, it does not always do so. In the rapid-clearing model of osteosarcoma, CAR T cells efficiently clear the cancer and do not elicit neuroinflammatory changes of CSF cytokine/chemokine elevation nor microglial reactivity. This suggests that it is not the CAR T cell interaction with the tumor per se, but rather the broader immune response frequently evoked by anti-tumor CAR T responses that results in these neuropathophysiological changes. While much remains to be learned about how the immune response evoked by tumor-clearing CAR T cell therapy outside of the CNS results in CNS inflammation, a similar phenomenon is observed after certain viral respiratory infections. Infection with SARS-CoV2 or H1N1 influenza that is restricted to the respiratory system also induces CSF cytokine and chemokine elevations and white matter-selective microglial reactivity^15^.

The cognitive deficits that can occur following chemotherapy, immunotherapy, and certain infections such as COVID or influenza share clinical features and neurobiological underpinnings. Microglia in the subcortical white matter, also called axon tract microglia^14^, are exquisitely sensitive to immune challenges and toxic exposures and exhibit reactivity following systemic methotrexate chemotherapy^11,12^, mild respiratory COVID^15^, mild respiratory H1N1 influenza^15^, and CAR T cell therapy. In mild respiratory-restricted COVID or influenza mouse models^15^, and in the aggressive osteosarcoma model studied here, neuroinflammatory changes are seen in response to immune challenges exclusively outside of the nervous system. How the immune response is relayed from the periphery to the CNS is incompletely understood. Past work demonstrates that circulating chemokines such as CCL11 are sufficient to elicit microglial reactivity and impaired neurogenesis^15,29^, but the possible signaling molecules, immune cellular trafficking and/or afferent neural input responsible for the broader neuroinflammatory response remain to be elucidated. However, a range of disease states characterized by a syndrome of impaired attention, memory, speed of information processing and executive function share a consistent pattern of white matter microglial reactivity, multi-lineage cellular dysregulation disrupting neural homeostasis and plasticity, and consequent cognitive impairment. Neural-immune principles of “brain fog” syndromes are thus emerging, and therapeutic strategies developed to mitigate cognitive symptoms for one disease context are likely to apply to many.

While consistent neuropathophysiological patterns are emerging in the various disease contexts associated with “brain fog” syndromes, important distinctions are also emerging. Comparing myelin changes after respiratory COVID, methotrexate chemotherapy and CAR T cell therapy, myelinated axons are decreased in each case, but only after methotrexate is myelin sheath thickness also affected. Comparing respiratory infections with SARS-CoV2 or H1N1 influenza, white matter microglial reactivity and oligodendroglial dysregulation were evident at 7 days in both and resolved by 7 weeks after influenza but remained aberrant at 7 weeks after SARS-CoV2 infection. Comparing cytokine and chemokine profiles in the CSF after these two respiratory viral infections reveals both shared and distinct patterns of persistent elevation. Similarly, comparing CSF cytokine/chemokine elevations in aggressive osteosarcoma and ALL models following tumor-clearing CAR T cell therapy here, many shared but also some distinct changes are observed, such as elevated CCL7 and BAFF in the ALL model but not the aggressive osteosarcoma model. Finally, important distinctions between subpopulations of microglia are important to recognize. While white matter microglia are uniquely sensitive to various toxic and immune challenges, microglia in hippocampal white matter are further distinguished by differential sensitivity to CCL11^15^ and resistance to CSF1Ri-mediated clearance.

## Limitations of the Study

An important caveat of this study is the use of immunocompromised mouse models. While this enabled evaluation of the CAR T cells used in the clinic and patient-derived tumors, these mice lack endogenous lymphocytes, and while myeloid cells are present these are not educated by lymphocytes as would normally occur in an immunocompetent mouse^45^. The full immune response elicited by tumor-clearing CAR T cell therapy may therefore not be appreciated here. Another important caveat of studying patient-derived xenograft models treated with human CAR T cells is that graft-versus-host disease develops after approximately 6 weeks, so longer term effects on cognition could not be studied here. A remaining question is thus how long the cellular dysregulation and cognitive impairment may persist beyond the timepoints assessed here.

Additional mechanisms implicated in cancer therapy-related cognitive impairment, and future studies should explore the role these mechanisms may play in ICRI. In addition to oligodendroglial and myelin dysregulation, reactive microglia can also induce neurotoxic substates of astrocytes^16,46^, which may cause cell death of oligodendrocytes and susceptible neurons, as well as impair normal astrocytic support of synapses^16^. Such astrocyte reactivity has been described following methotrexate chemotherapy in mice, and might explain the loss of oligodendroglial homeostasis observed here after CAR T cell therapy. Impaired synaptic connectivity and plasticity can be another consequence of traditional cancer therapies^47–49^, and should be explored in future studies of IRCI. Finally, chemotherapy-induced choroid plexus dysfunction may alter the antioxidant properties of CSF and contribute to cognitive impairment ^50^. Whether similar such alterations in choroid plexus biology and consequent changes to CSF supportive and neurotrophic functions^51^ are induced by immunotherapy remains to be determined.

## Conclusions

CAR T cell therapy has been transformative for hematological malignancies and holds enormous promise for solid tumors. Understanding the potential cognitive sequelae of the immune response associated with tumor-clearing CAR T-cell therapy and the neuroimmunological underpinnings of these possible neurological consequences is of paramount importance to optimizing quality of life for survivors. Given the mechanistic parallels evident in cognitive impairment syndromes that follow a range of inflammatory disease states, such progress may enable therapeutic interventions for many living with persistent “brain fog” syndromes.

## Acknowledgments

This work was supported by grants from the National Institute of Neurological Disorders and Stroke (R01NS092597), NIH Director’s Pioneer Award (DP1NS111132), National Cancer Institute (P50CA165962, R01CA258384, U19CA264504), Cancer Research UK, Robert J. Kleberg, Jr. and Helen C. Kleberg Foundation, McKenna Claire Foundation, Unravel Pediatric Cancer Foundation, ChadTough Defeat DIPG, Alex’s Lemonade Stand Foundation, Kyle O’Connell Foundation, Virginia and D.K. Ludwig Fund for Cancer Research, Waxman Family Research Fund, Will Irwin Research Fund, Parekh Center for Interdisciplinary Neurology, National Eye Institute R01EY033353, MD Anderson Cancer Center Neurodegeneration Consortium. The authors thank the Human Immune Monitoring Core at Stanford. Model schematics and graphical abstract were designed with Biorender.com

## Author Contributions

Conceptualization, methodology, validation, and visualization were performed by M.M., R.G.M., A.C.G., L.A-A.; formal analysis by A.C.G., L.A-A., M.O’D; resources provided by M.M., R.G.M.; investigation was performed by A.C.G., L.A-A., M.R., S.D., R.M., K.S.; writing – original draft by M.M. and L.A-A; writing – review & editing by M.M., A.C.G., L.A-A, K.S.; data curation by A.C.G., L.A-A.; supervised by M.M., S.A.L., R.G.M.; funding acquisition by M.M., S.A.L., R.G.M

---

**Supplemental Figure 1:**
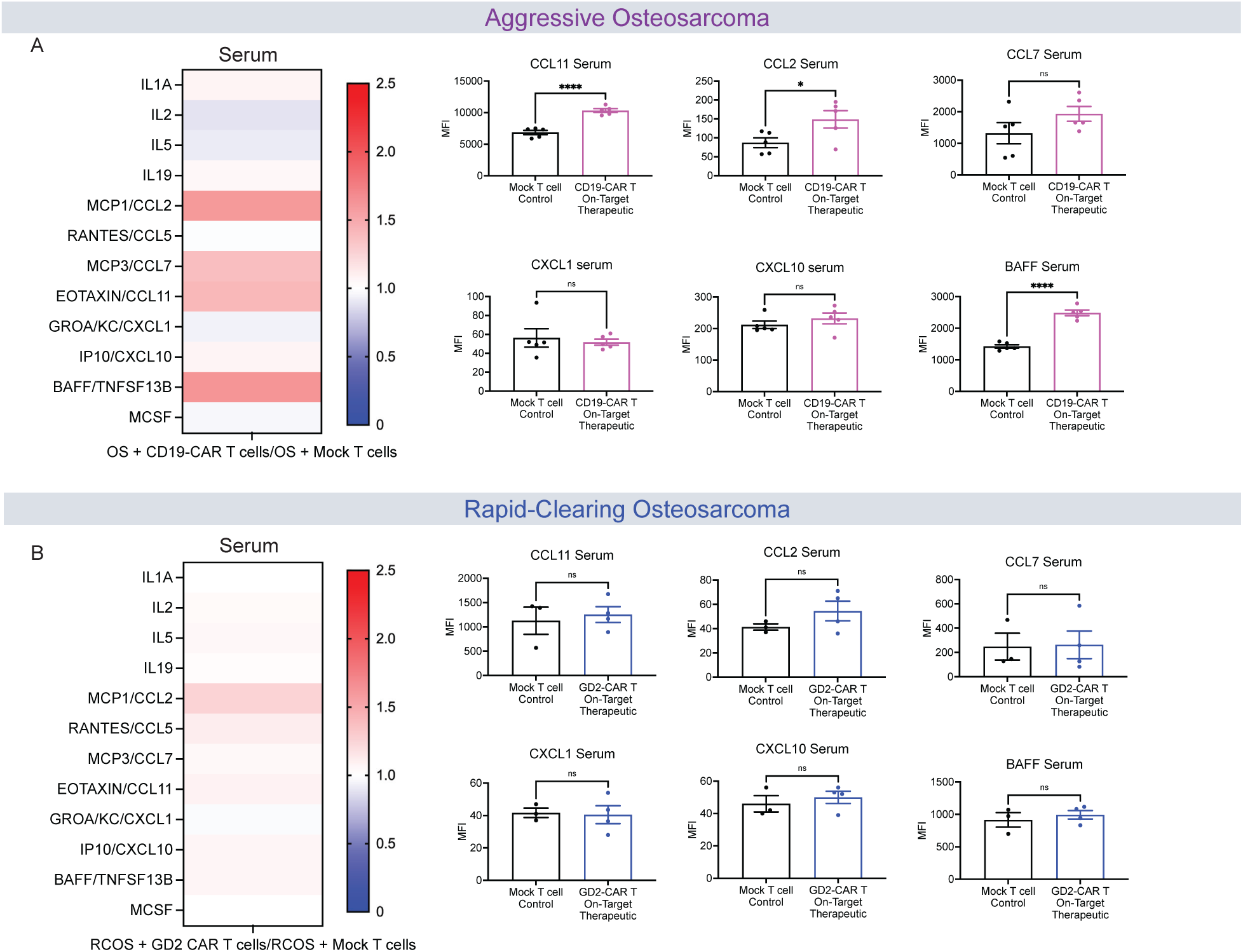
Cytokine/chemokine measurements in the serum in two Osteosarcoma models. (A) Left, heat map of fold change of cytokine and chemokine analysis of serum in aggressive osteosarcoma (OS)+On-Target Therapeutic CD19-CAR T cell treated mice relative to aggressive osteosarcoma (OS)+mock T cell treated control mice 35-days post-CAR T cell therapy. Right, quantification of raw MFI of serum levels of cytokines (CCL11, CCL2, CCL7, CXCL1, CXCL10, BAFF). (B) Left, heat map of fold change of cytokine and chemokine analysis of serum in rapid-clearing osteosarcoma (RCOS)+On-Target Therapeutic GD2-CAR T cell treated mice relative to rapid-clearing osteosarcoma (RCOS)+mock T cell treated control mice 35-days post-CAR T cell therapy. Right, quantification of raw MFI of serum levels of cytokines (CCL11, CCL2, CCL7, CXCL1, CXCL10, BAFF). (A and B) Cytokine and chemokines significantly elevated or depressed, analyzed via Student’s T test of raw MFI values.

**Supplemental Figure 2.**
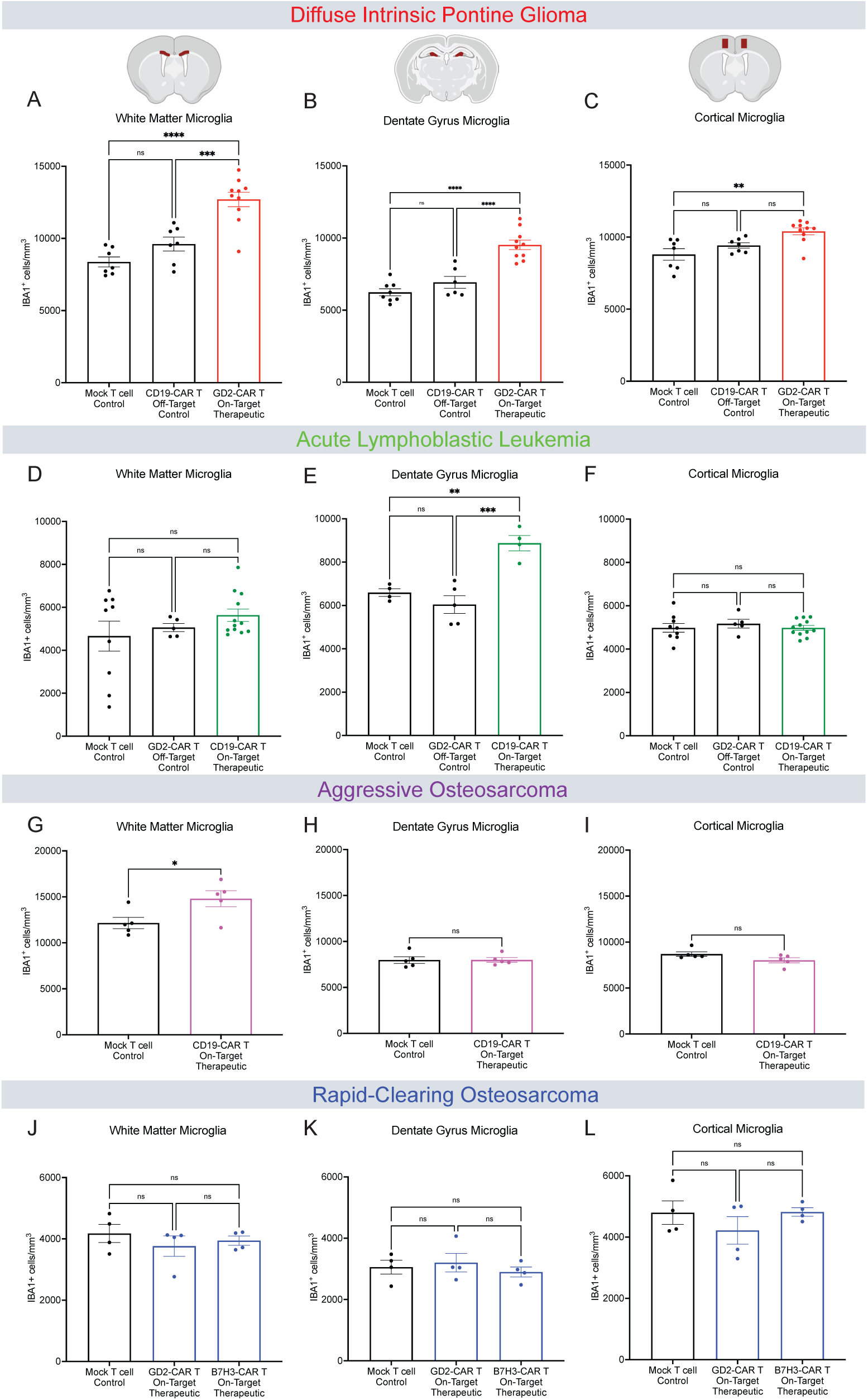
Microglial density in the white matter, dentate gyrus, and cortex. (A) Microglia (IBA1+) quantification in the corpus callosum white matter 35 days post-CAR T therapy in the DIPG model. Mock T cell Control (n=7 mice), CD19-CAR T Control (n=7 mice), GD2-CAR T Therapeutic (n=10 mice). (B) Microglia (IBA1+) quantification in the hilar white matter region of the dentate gyrus 35 days post-CAR T therapy in the DIPG model. Mock T cell Control (n=8 mice), CD19-CAR T Control (n=6 mice), GD2-CAR T Therapeutic (n=10 mice). (C) Microglia (IBA1+) quantification in the cortical gray matter 35 days post-CAR T therapy in the DIPG model. Mock T cell Control (n=7 mice), CD19-CAR T Control (n=7 mice), GD2-CAR T Therapeutic (n=10 mice). (D) Microglia (IBA1+) quantification in the corpus callosum white matter 35 days post-CAR T therapy in the ALL model. Mock T cell Control (n=9 mice), GD2-CAR T Control (n=5 mice), CD19-CAR T Therapeutic (n=12 mice). (E) Microglia (IBA1+) quantification in the hilar white matter region of the dentate gyrus 35 days post-CAR T therapy in the ALL model. Mock T cell Control (n=4 mice), GD2-CAR T Control (n=5 mice), CD19-CAR T Therapeutic (n=4 mice). (F) Microglia (IBA1+) quantification in the cortical gray matter 35 days post-CAR T therapy in the ALL model. Mock T cell Control (n=9 mice), GD2-CAR T Control (n=5 mice), CD19-CAR T Therapeutic (n=12 mice). (G) Microglia (IBA1+) quantification in the corpus callosum white matter 35 days post-CAR T therapy in the aggressive osteosarcoma model. Mock T cell Control (n=5 mice), CD19-CAR T Therapeutic (n=5 mice). (H) Microglia (IBA1+) quantification in the hilar white matter region of the dentate gyrus 35 days post-CAR T therapy in the aggressive osteosarcoma model. Mock T cell Control (n=5 mice), CD19-CAR T Therapeutic (n=5 mice). (I) Microglia (IBA1+) quantification in the cortical gray matter 35 days post-CAR T therapy in the aggressive osteosarcoma model. Mock T cell Control (n=5 mice), CD19-CAR T Therapeutic (n=5 mice). (J) Microglia (IBA1+) quantification in the corpus callosum white matter 35 days post-CAR T therapy in the rapid-clearing osteosarcoma model. Mock T cell Control (n=4 mice), GD2-CAR T Therapeutic (n=4 mice), B7H3-CAR T Therapeutic (n=4 mice). (K) Microglia (IBA1+) quantification in the hilar white matter region of the dentate gyrus 35 days post-CAR T therapy in the rapid-clearing osteosarcoma model. Mock T cell Control (n=4 mice), GD2-CAR T Therapeutic (n=4 mice), B7H3-CAR T Therapeutic (n=4 mice). (L) Microglia (IBA1+) quantification in the cortical gray matter 35 days post-CAR T therapy in the rapid-clearing osteosarcoma model. Mock T cell Control (n=4 mice), GD2-CAR T Therapeutic (n=4 mice), B7H3-CAR T Therapeutic (n=4 mice). Data shown as mean ± SEM (A-F, J-L). Each point = one mouse. ns = p > 0.05, *p < 0.05, **p < 0.01, ***p < 0.001, ****p < 0.0001, analyzed via One-way ANOVA. (G-I) Data shown as mean ± SEM. Each point = one mouse, ns = p > 0.05, *p < 0.05, analyzed via Unpaired T Test.

**Supplemental Figure 3.**
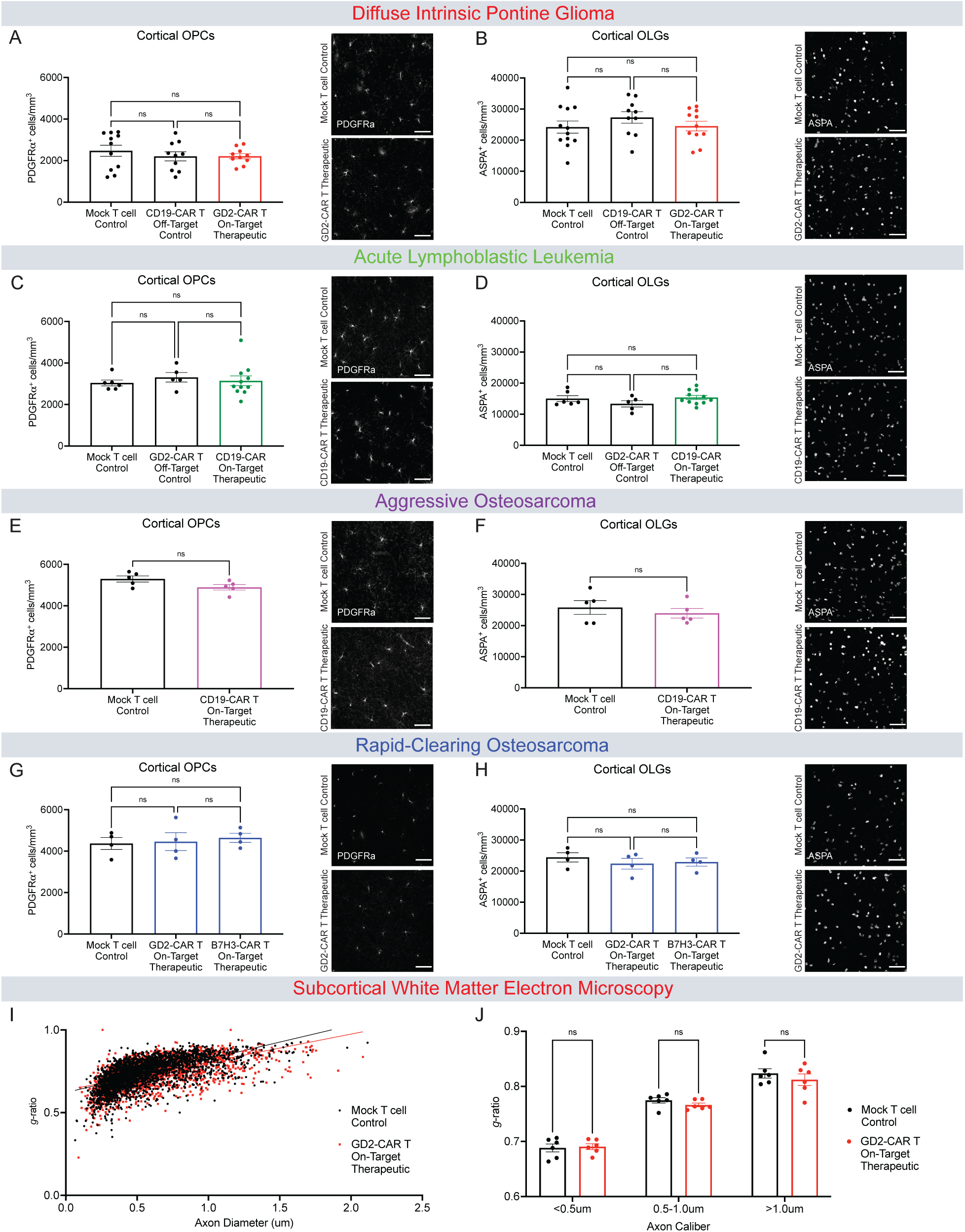
Normal cortical oligodendroglia and subcortical myelin sheath thickness following tumor-clearing CAR T cell therapy. (A) Quantification and representative confocal micrographs of PDGFRa+ oligodendrocyte precursor cells from the cortical gray matter in DIPG xenografted mice 35 days post-CAR T therapy. Mock T cell Control (n=11 mice), CD19-CAR T Control (n=10 mice), GD2-CAR T Therapeutic (n=10 mice). (B) Quantification and representative confocal micrographs of ASPA+ mature oligodendrocytes from the cortical gray matter in DIPG xenografted mice 35 days post-CAR T therapy. Mock T cell Control (n=12 mice), CD19-CAR T Control (n=10 mice), GD2-CAR T Therapeutic (n=11 mice). (C) Quantification and representative confocal micrographs of PDGFRa+ oligodendrocyte precursor cells from the cortical gray matter in ALL xenografted mice 28 days post-CAR T therapy. Mock T cell Control (n=6 mice), GD2-CAR T Control (n=5 mice), CD19-CAR T Therapeutic (n=11 mice). (D) Quantification and representative confocal micrographs of ASPA+ mature oligodendrocytes from the cortical gray matter in ALL xenografted mice 28 days post-CAR T therapy. Mock T cell Control (n=6 mice), CD19-CAR T Control (n=5 mice), GD2-CAR T Therapeutic (n=10 mice). (E) Quantification and representative confocal micrographs of PDGFRa+ oligodendrocyte precursor cells from the cortical gray matter in aggressive osteosarcoma xenografted mice 35 days post-CAR T therapy. Mock T cell Control (n=5 mice), CD19-CAR T Therapeutic (n=5 mice). (F) Quantification and representative confocal micrographs of ASPA+ mature oligodendrocytes from the cortical gray matter in aggressive osteosarcoma xenografted mice 35 days post-CAR T therapy. Mock T cell Control (n=5 mice), CD19-CAR T Therapeutic (n=5 mice). (G) Quantification and representative confocal micrographs of PDGFRa+ oligodendrocyte precursor cells from the cortical gray matter in rapid-clearing osteosarcoma xenografted mice 35 days post-CAR T therapy. Mock T cell Control (n=4 mice), GD2-CAR T Therapeutic l (n=4 mice), B7H3-CAR T Therapeutic (n=4 mice). (H) Quantification and representative confocal micrographs of ASPA+ mature oligodendrocytes from the cortical gray matter in rapid-clearing osteosarcoma xenografted mice 35 days post-CAR T therapy. Mock T cell Control (n=4 mice), GD2-CAR T Therapeutic l (n=4 mice), B7H3-CAR T Therapeutic (n=4 mice). (I) Scatterplots of g-ratio relative to axon diameter at 35 days post-CAR T cell therapy for mock T cell control (black points) or GD2-CAR T On-Target Therapeutic (red points) treated mice. (n=6 mice per group) (J) G-ratio relative to small (<0.5um), medium (0.5-1.0um), and large (>1.0um) caliber axons for mock T cell control (black points) or GD2-CAR T On-Target Therapeutic (red points) treated mice. (n=6 mice per group) (A-D, G-H, J) Data shown as mean ± SEM. Each point = one mouse. ns = p > 0.05, analyzed via One-way ANOVA. (E & F) Data shown as mean ± SEM. Each point = one mouse, ns = p > 0.05, analyzed via Unpaired T Test. Scale bars equal 50um in all confocal images.

**Supplemental Figure 4.**
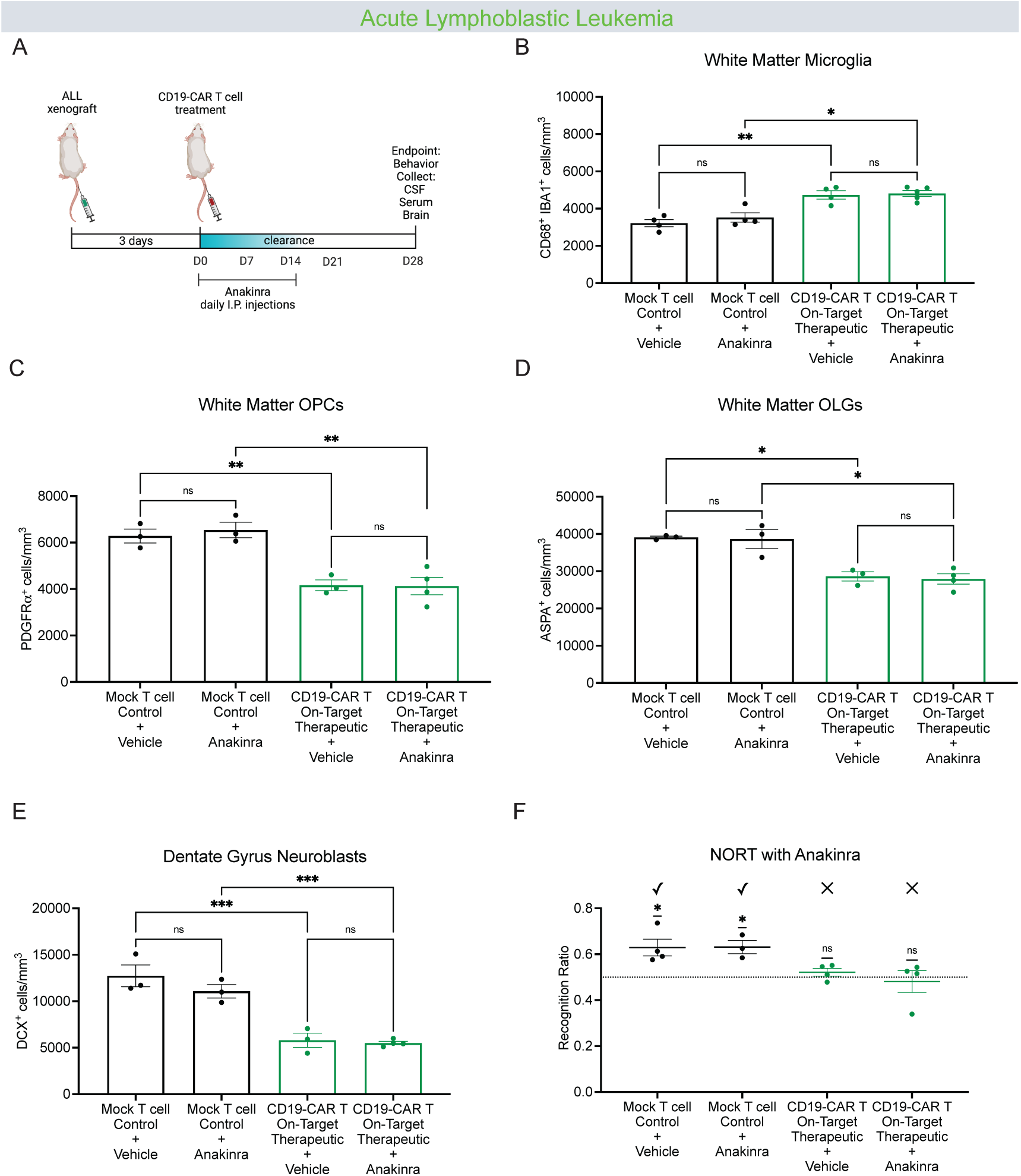
Anakinra does not rescue cellular or behavioral deficits. (A) Schematic illustration denoting the timeline for the ALL model with Anakinra administration. (B) Quantification of corpus callosum reactive white matter microglia (IBA1+ CD68+) 28 days post-CAR T therapy with or without 14 days of Anakinra administration in ALL-xenografted mice. Mock T cell+Vehicle (n=4 mice), mock T cell+Anakinra (n=4 mice), CD19-CAR T Therapeutic+Vehicle (n=4 mice), CD19-CAR T Therapeutic+Anakinra (n=5 mice). (C) Quantification of PDGFRa+ oligodendrocyte precursor cells from the corpus callosum in ALL-xenografted mice 28 days post-CAR T therapy with or without 14 days of Anakinra administration. Mock T cell+Vehicle (n=4 mice), mock T cell+Anakinra (n=4 mice), CD19-CAR T Therapeutic+Vehicle (n=4 mice), CD19-CAR T Therapeutic+Anakinra (n=5 mice). (D) Quantification of ASPA+ mature oligodendrocytes from the corpus callosum in ALL-xenografted mice 28 days 28 days post-CAR T therapy with or without 14 days of Anakinra administration. Mock T cell+Vehicle (n=4 mice), mock T cell+Anakinra (n=4 mice), CD19-CAR T Therapeutic+Vehicle (n=4 mice), CD19-CAR T Therapeutic+Anakinra (n=5 mice). (E) Quantification of dentate gyrus neuroblasts (DCX+) 28 days post-CAR T therapy with or without 14 days of Anakinra administration in ALL-xenografted mice. Mock T cell+Vehicle (n=3 mice), mock T cell+Anakinra (n=3 mice), CD19-CAR T Therapeutic+Vehicle (n=3 mice), CD19-CAR T Therapeutic+Anakinra (n=4 mice). (F) NORT performed for ALL-xenografted mice analyzed on day 28 post-CAR T therapy with or without Anakinra administration. Mock T cell+Vehicle (n=4 mice), mock T cell+Anakinra (n=3 mice), CD19-CAR T Therapeutic+Vehicle (n=4 mice), CD19-CAR T Therapeutic+Anakinra (n=4 mice). (B, C, D, E) Data shown as mean ± SEM. Each point = one mouse. ns = p > 0.05, **p < 0.01, ***p < 0.001, analyzed via 2-way ANOVA. (F) Data shown as mean ± SEM. Each point = one mouse. ns = p > 0.05, *p < 0.05, **p < 0.01, ****p < 0.0001. Analyzed via One Sample T and Wilcoxon Test.

**Supplemental Figure 5.**
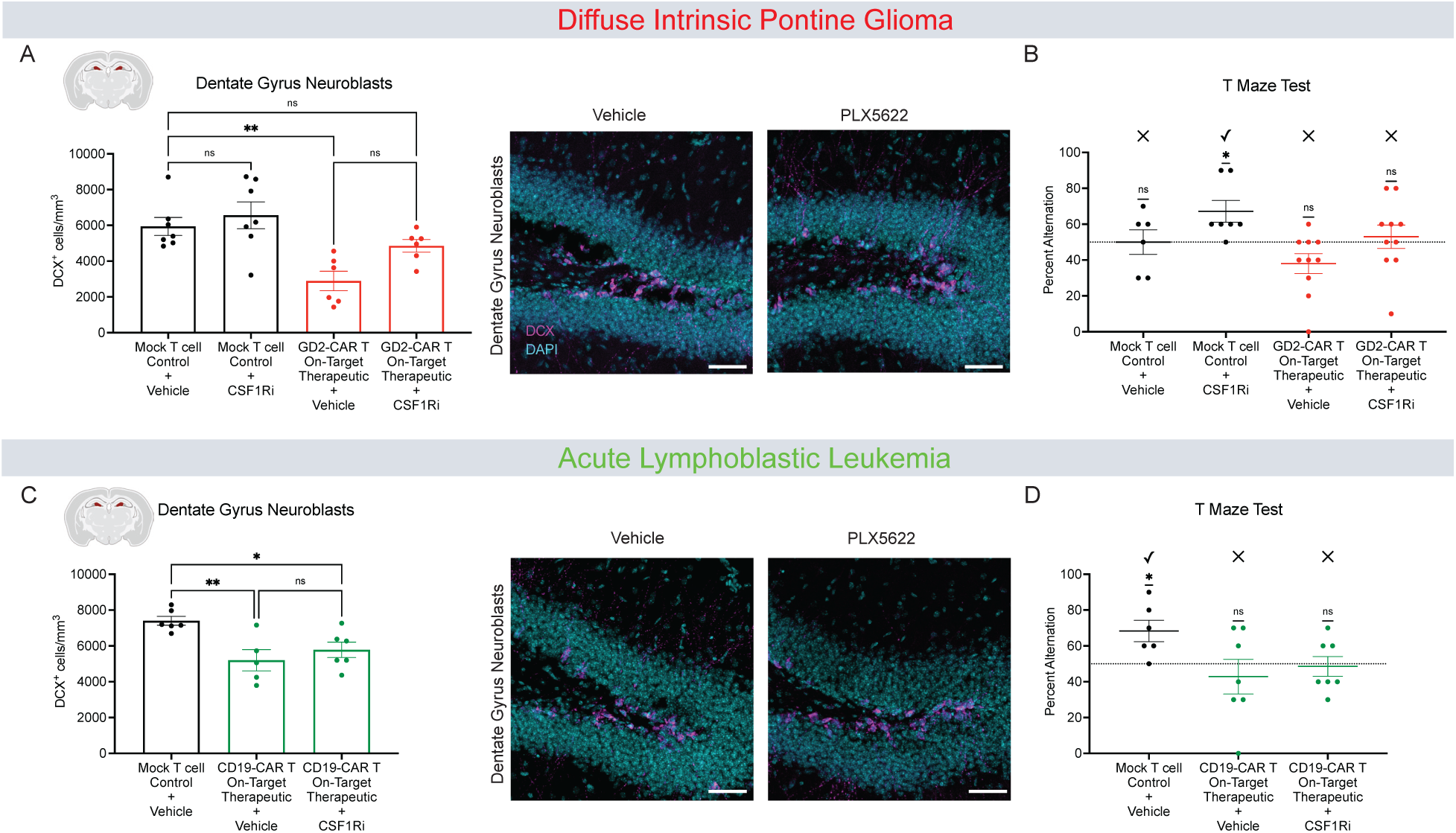
Incomplete microglial depletion in the hippocampus does not rescue behavioral deficits. (A) Quantification of dentate gyrus neuroblasts (DCX+) 35 days post-CAR T therapy with or without 14 days of CSF1R inhibition in DIPG-xenografted mice,. Mock T cell+Vehicle Chow (n=7 mice), mock T cell+CSF1Ri Chow (n=7 mice), GD2-CAR T Therapeutic+Vehicle Chow (n=6 mice), GD2-CAR T Therapeutic+CSF1Ri Chow (n=6 mice). Representative confocal micrographs of neuroblasts (magenta = DCX, cyan = DAPI) in the dentate gyrus of DIPG xenografted mice 35 days after CAR T cell therapy with or without 14 days of CSF1R inhibitor (PLX5622). (B) T Maze Test results from DIPG-xenografted mice D35 post-CAR T therapy with or without 14 days of CSF1R inhibition. Mock T cell+Vehicle Chow (n=6 mice), mock T cell+CSF1Ri Chow (n=7 mice), GD2-CAR T Therapeutic+Vehicle Chow (n=10 mice), GD2-CAR T Therapeutic+CSF1Ri Chow (n=10 mice). (C) Quantification of dentate gyrus neuroblasts (DCX+) in ALL-xenografted mice at 28 days post-CAR T therapy with or without 14 days of CSF1R inhibition. Mock T cell+Vehicle Chow (n=6 mice), CD19-CAR T Therapeutic+Vehicle Chow (n=5 mice), CD19-CAR T Therapeutic+CSF1Ri Chow (n=6 mice). Representative confocal micrographs of neuroblasts (magenta = DCX, cyan = DAPI) in the dentate gyrus of ALL xenografted mice 28 days after CAR T cell therapy with or without 14 days of CSF1R inhibitor (PLX5622). (D) T maze Test results from ALL-xenografted mice at D28 post-CAR T therapy with or without 14 days of CSF1R inhibition. Mock T cell+Vehicle Chow (n=6 mice), CD19-CAR T Therapeutic+Vehicle Chow (n=7 mice), CD19-CAR T Therapeutic+CSF1Ri Chow (n=7 mice). Data shown as mean ± SEM (A-D). Each point = one mouse. ns = p > 0.05, * p < 0.05, **p < 0.01, analyzed via 2-way ANOVA (A, C), One Sample T and Wilcoxon Test (B, D). Scale bars equal 50um in all confocal images.

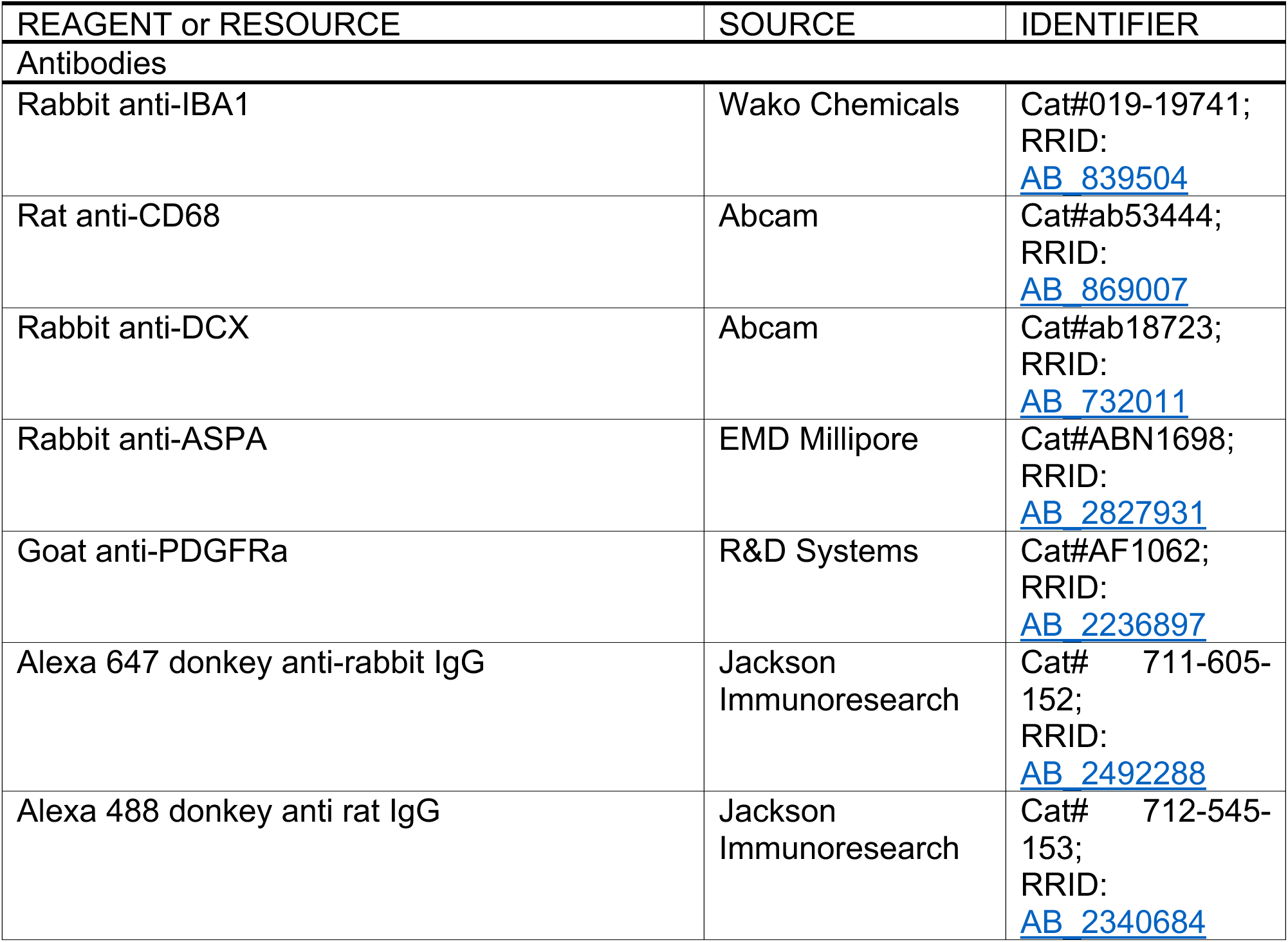
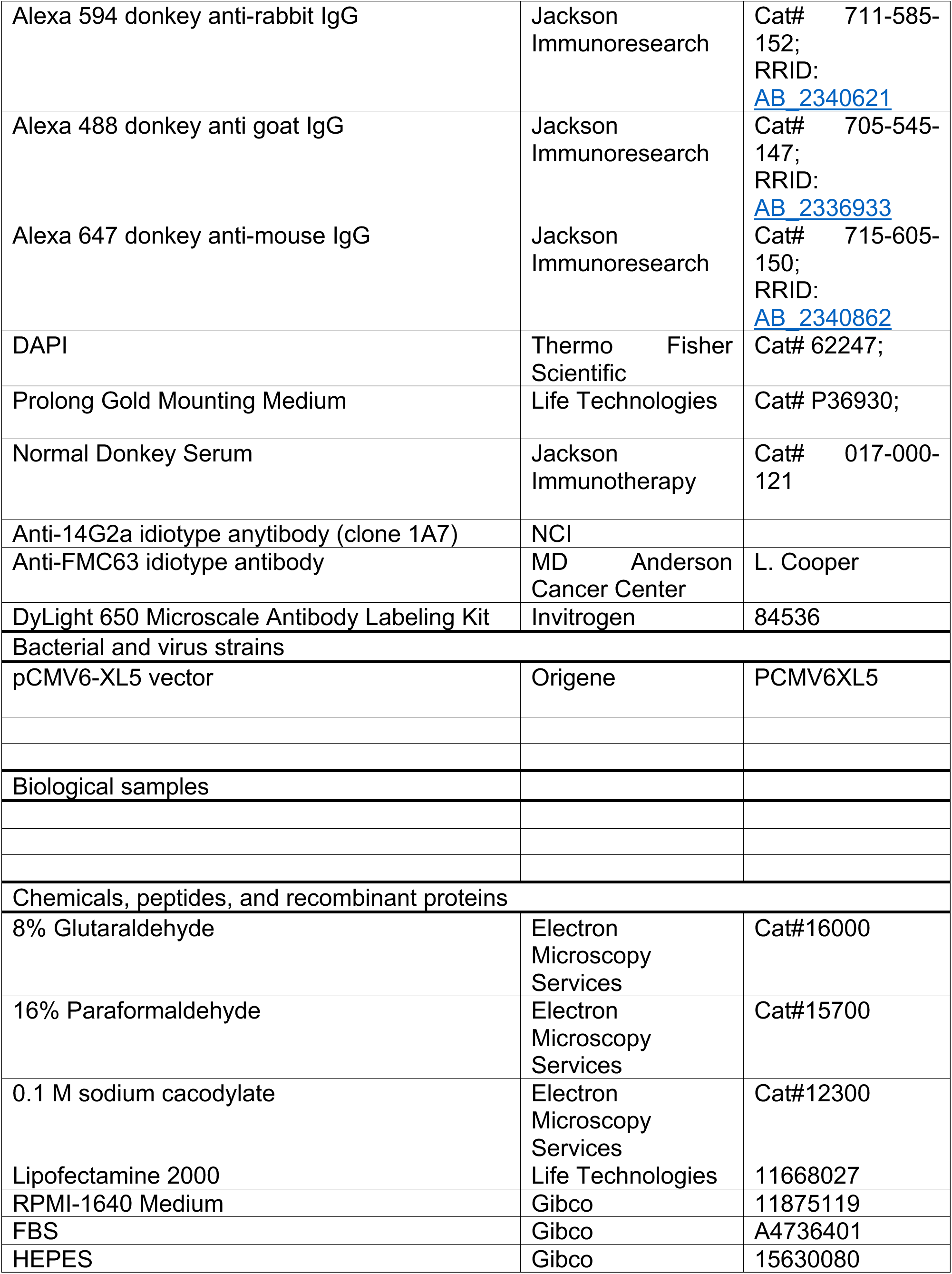
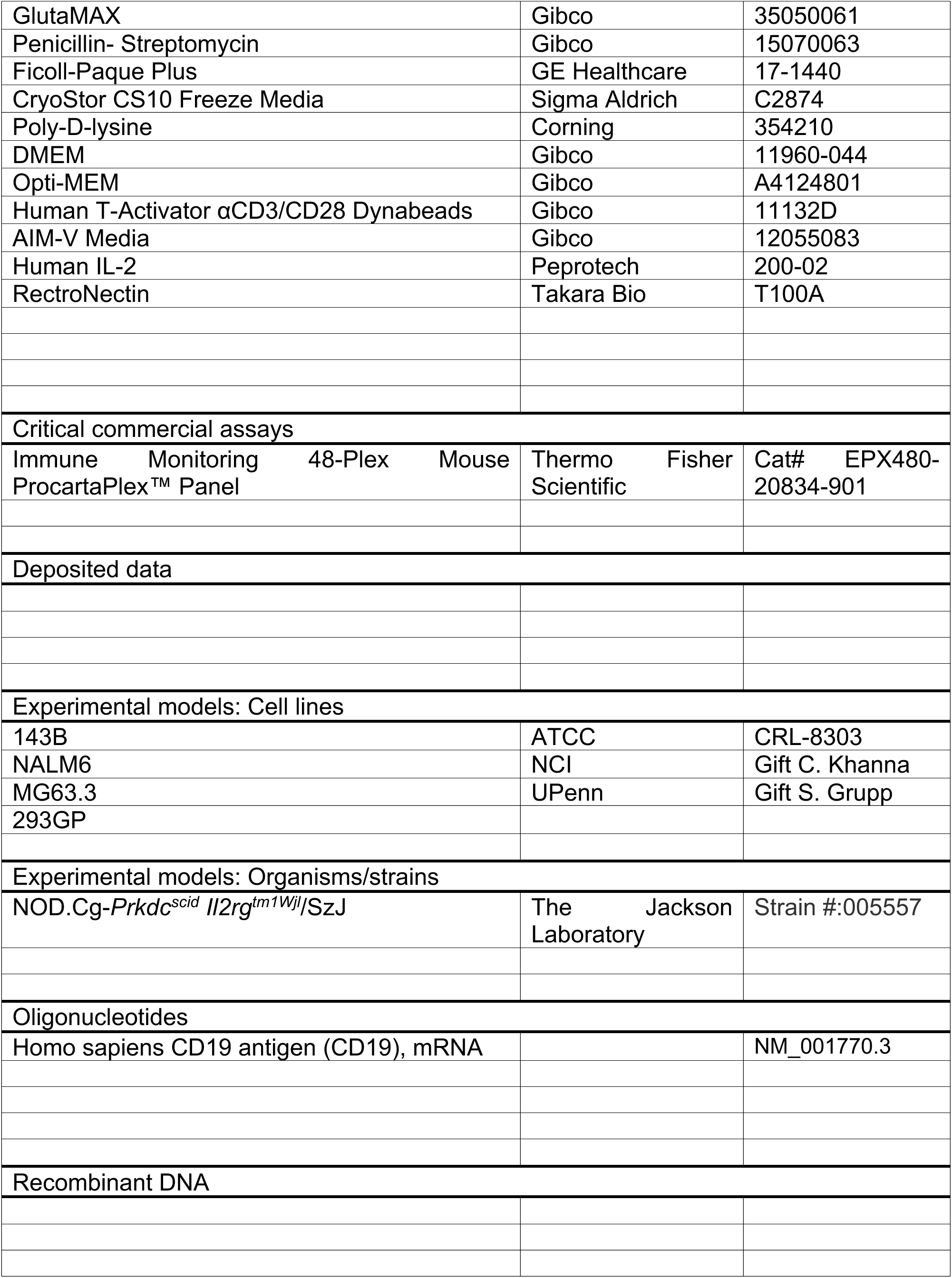
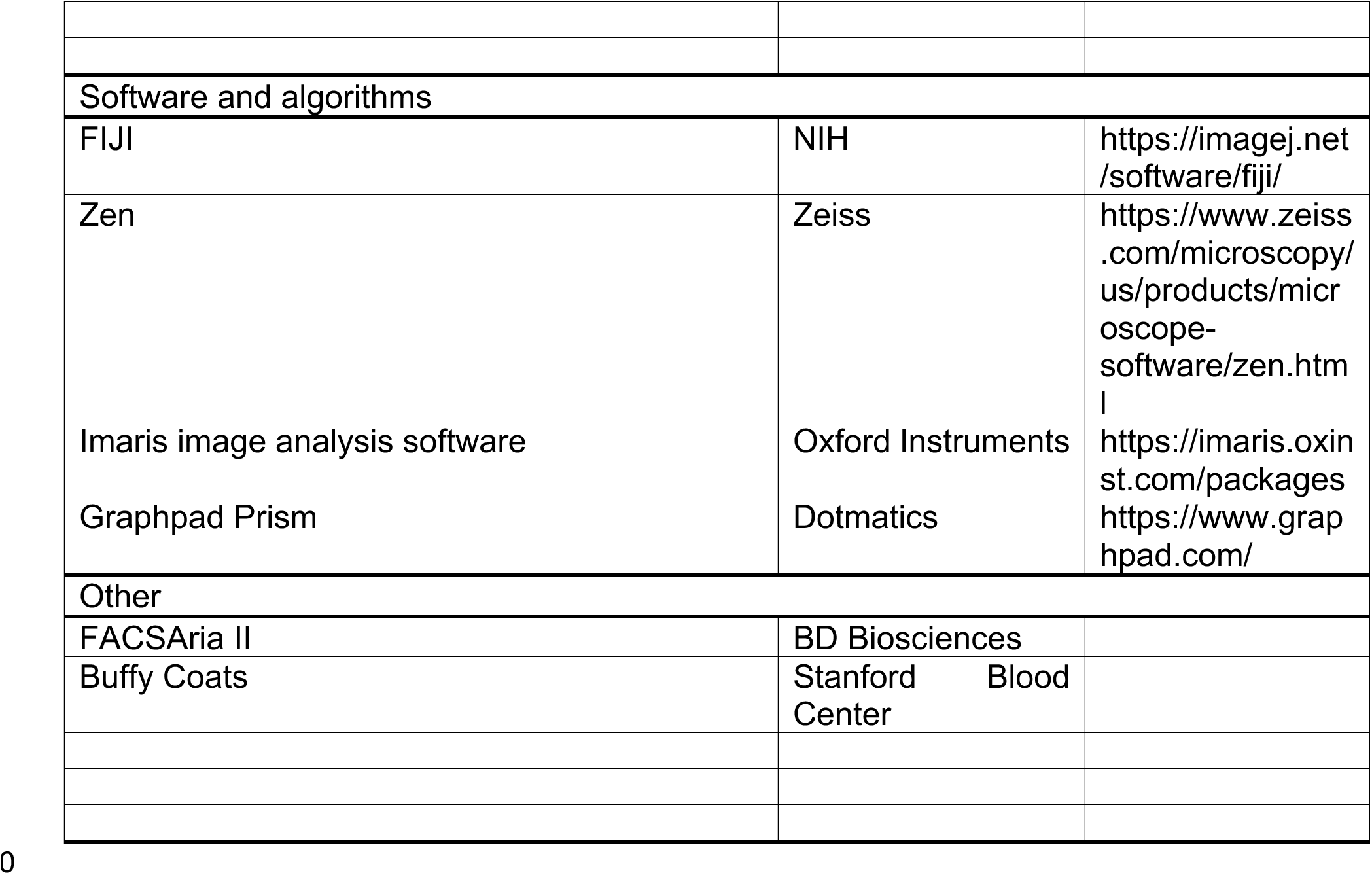
KEY RESOURCES TABLE

## STAR Methods

### RESOURCE AVAILABILITY

#### Lead Contact

Further information and requests for resources and reagents should be directed to and will be fulfilled by Michelle Monje

### Materials Availability

### EXPERIMENTAL METHOD AND SUBJECT DETAILS

#### Cell and Culture conditions

The Nalm6-GFP Luciferase B-ALL cell line was obtained from S. Grupp (University of Pennsylvania, Philadelphia, PA). The 143B cell line was obtained from ATCC and was transfected with CD19 (NM_001770.3) in the pCMV6-XL5 vector (Origene) using Lipofectamine 2000 (Life Technologies). Transfected cells have been sorted (FACSAria II, BD Bioscience) and single cell cloned to obtain a pure population ^24^. MG63.3 was obtained by C. Khanna (NCI, NIH, Bethesda, MD). All the cell lines were cultured in RPMI-1640 medium (Gibco) supplemented with 10% FBS, 10 mM HEPES, 2 mM GlutaMAX, 100 U/mL penicillin, and 100 μg/mL streptomycin (Gibco).

#### PBMC isolation

Buffy coats, leukopaks or Leukocyte Reduction System (LRS) chambers from healthy donor were obtained through the Stanford Blood Center under an IRB-exempt protocol. Peripheral blood mononuclear cells were isolated using Ficoll-Paque Plus (GE Healthcare, 17-1440) density gradient centrifugation according to the manufacturer’s instructions and cryopreserved with CryoStor CS10 freeze media (Sigma-Aldrich) in 2-5 x 10^7^ cell aliquots.

#### CAR construction

The GD2-4-1-BBz and CD19-4-1-BBz CARs have been previously described ^24^and were generated by cloning the scFvs derived from the 14G2a and FMC63 antibodies into the MSGV1 retroviral vector. In some experiments, CAR vectors including the mcherry reporter gene after the CD3z signaling domain were used.

#### Virus production

Retroviral supernatant was generated as previously described ^24^. Briefly, 6-7 M of 293GP cells were plated on 100mm poly-D-lysine (Corning) coated plates in complete DMEM media supplemented with 10% FBS, 10 mM HEPES, 2 mM GlutaMAX, 100 U/mL penicillin, and 100 μg/mL streptomycin (Gibco). The day after, cells were transfected with 4.5 μg of RD114 and 9 μg of the CAR plasmid using Lipofectamine 2000 (Invitrogen) in Opti-MEM media (Gibco). Media was changed 24 hours after transfection. The supernatant was harvested after 48 and 72 hours and stored at -80°C until use.

#### CAR T generation

CAR T cells were generated as previously described ^52^ (Walker, A. J. et al. Tumor Antigen and Receptor Densities Regulate Efficacy of a Chimeric Antigen Receptor Targeting Anaplastic Lymphoma Kinase. Molecular Therapy 25, 2189-2201, doi:10.1016/j.ymthe.2017.06.008). Briefly, at day 0, PBMCs were thawed and activated with Human T-Activator αCD3/CD28 Dynabeads (Gibco) at a 3:1 bead to cell ratio, in AIM-V media (Gibco) supplemented with 5% FBS, 10 mM HEPES, 2 mM GlutaMAX, 100 U/mL penicillin, 100 μg/mL streptomycin, and 100 U/mL recombinant human IL-2 (Peprotech). Retroviral transduction was done on day 3 and 4 by using non-tissue culture treated plates previously coated with RetroNectin (Takara Bio) according to manufacturer’s instructions. One mL of retroviral supernatant was added and the plates were then centrifuged at 3,200 RPM at 32°C for 2 hours. The supernatant was then removed and 0.5 x 106 T cells were added to each well in 1 mL of complete AIM-V media. The Dynabeads were removed on day 5 by magnetic separation and the T cells were maintained in culture at a density between 0.3 – 0.5×106 cells/mL until Day 10, with media changes every 2-3 days.

#### Flow cytometry

GD2 CAR expression was verified by using the anti-14G2a idiotype antibody (clone 1A7) obtained from National Cancer Institute^53^, together with mCherry detection, when present. CD19 CAR expression was detected with an anti-FMC63 idiotype antibody provided by L. Cooper (MD Anderson Cancer Center)^54^. Idiotype antibodies were fluorescently labeled with the DyLight 650 Microscale Antibody Labeling Kit (Invitrogen). The cells were washed and stained in FACS buffer (PBS supplemented with 2% FBS) for 20 minutes at 4°C. Flow cytometry was performed on a BD Fortessa instrument.

#### Mouse models

All studies were conducted with NSG mice (NOD-SCID-IL2R gamma chain-deficient, Jackson Laboratories). Males and females were used equally, and all animals were housed at Stanford University; all procedures complied with Institutional Animal Care and Use Committee protocol at Stanford. Animals were housed in a standard 12h light: 12h dark cycle with ad libitum access to food and water.

#### Diffuse Intrinsic Pontine Glioma *in vivo* model

For the orthotopic xenograft DIPG model, NSG mice were bred at Stanford following all approved protocols. Pups between the ages of P0-P3 were anesthetized by cryoanesthesia. Mice were placed on a piece of sterile gauze on ice until they are no longer moving. Then mice were stabilized on a stereotactic surgical frame (Stoelting) cooled down to ensure continuous deep cryoanesthesia of the pups. A single cell suspension of SU-DIPG-VI-GFP was prepared in sterile HBSS immediately before injection. Approximately 100,000 cells in 2ul of HBSS was injected at a rate of 0.5 microliter per minute using a 33 gauge Hamilton needle into the fourth ventricle of the brain at the following coordinates: 3 mm dorsal to bregma and 3 mm deep to cranial surface. The needle remained in the brain for one minute before retraction and animals were monitored after surgery while they recovered on a heating pad. Mice were then returned to their mother and monitored until normal weaning at P21. Eight weeks after tumor injection, mice are assessed for tumor burden via in vivo luminescence imaging (IVIS Spectrum, Perkin Elmer) and total flux was calculated using the machine’s software (Living Image, Perkin Elmer) measured as circular ROIs centered over the mouse’s skull. Mice were ranked by tumor burden and then sequentially distributed over three different groups: GD2-4-1-BBz CAR T-cells (therapeutic), CD19-4-1-BBz CAR T-cells (non-targeting controls), or mock T-cells (activated but not infected with a viral construct). Mice were administered 1×10^7^ transduced cells in 200 microliters sterile HBSS. Tumor burden was measured weekly by IVIS to determine successful tumor clearance. Tumors were clear by D21 post injection, and then animals were left undisturbed in their home cages for two weeks after tumor clearance. On D35 post CAR T-cell injection, animals were subjected to behavioral assays and then CSF and brains were collected as described below.

#### Acute Lymphoblastic Leukemia *in vivo* model

Nalm6-GFP Luciferase B-ALL cells were cultured and expanded as described above and harvested with TrypLE Express (Gibco 12604-013). For the orthotopic model, 1×10^6^ cells in 200ul of sterile HBSS were injected intravenously in male and female mice aged 5-6 weeks. Three days after tumor injection, mice were assessed for tumor burden via IVIS, ranked by tumor burden and sequentially distributed over three groups: CD19-4BBz CAR T-cells (therapeutic), GD2-4BBz CAR T-cells (non-targeting controls), or mock T-cells (activated but not infected with a viral construct). Mice were administered 5×10^6^ transduced cells in 200ul of sterile HBSS and monitored twice a week by IVIS. To avoid infiltration of NALM6 cells to the brain, mock treated animals were assayed for cognitive differences and CSF and brain collection at D10 post NALM6 introduction, and brains were checked histologically for GFP+ cells. CAR T-cell treated animals cleared their tumors by D14 post CAR injection, and then were left undisturbed in their cages until D28, when they were assayed for cognition and tissue collection as described below.

#### Osteosarcoma *in vivo* models

MG63.3 and 143B-CD19 osteosarcoma lines were cultured and expanded as described above and harvested with TrypLE Express. For the orthotopic model, 1×10^6^ MG63.3 cells or 1×10^6^ 143B-CD19 cells were injected periosteal to the tibia of male and female mice aged 5-7 weeks. In the MG63.3 model, 4 weeks after tumor initiation animals were administered 5×10^6^ transduced GD2-4BBz CAR T-cells (therapeutic), B7H3 CAR T-cells (therapeutic) or mock T-cells. In the 143B-CD19 model, 1 week after tumor initiation animals were administered 10×10^6^ CD19-4BBz CAR T-cells (therapeutic) or mock T-cells. Tumors were measured with a digital caliper once a week. Two weeks after tumor clearance in each model (7D post treatment in MG63.3, 21D post treatment in 143B-CD19), animals were assayed for cognition and tissue collected as described below.

#### Anakinra treatment in the ALL model

Mice were engrafted with Nalm6-GFP Luciferase B-ALL cells as described above. Three days after tumor injection, mice were assessed for tumor burden via IVIS, ranked by tumor burden and sequentially distributed over four groups: CD19-4BBz CAR T-cells (therapeutic) +/-Anakinra, or mock T-cells +/-Anakinra. Mice were administered 5×10^6^ transduced cells in 200ul of sterile HBSS and monitored twice a week by IVIS. Beginning on Day 0 of CAR T cell treatment, animals were administered 10mg/kg of Kineret (Anakinra, Stanford Pharmacy) daily i.p., until Day 14 post treatment, the duration of tumor clearance. Mock treated animals were again measured for cognition at D10 post NALM6 engraftment, and CAR T-cell treated animals were left undisturbed in their cages until D28, when they were assayed for cognition and tissue collection as described below.

#### Microglia Depletion *in vivo* with PLX5622

To deplete microglia in the brain, mice in both the DIPG and ALL in vivo models were given *ad libitum* access to rodent chow containing 1200ppm of PLX5622 beginning after their tumors cleared (D21 post treatment in the DIPG model and D14 post treatment in the ALL model) and maintained on PLX5622 for two weeks. Animals underwent cognitive assays and CSF and brain collection as described below. PLX5622 was purchased from MedChemExpress (HY-114153) and formulated in AIN-76A standard chow by Research Diets Inc (D19101002i).

### METHOD DETAILS

#### CSF collection and perfusion

At the endpoint of each experiment as described above, mice were anesthetized with isoflurane and mounted onto a stereotactic surgical rig (Stoelting) with the head secured at a 30-degree angle facing upward. The hair of the neck was shaved and an incision was made through the skin and muscle. The muscle around the neck was separated with blunt metal forceps to reveal the cisterna magna. Any extra blood was cleaned with cotton tips until the area was cleared. CSF was removed from the cisterna magna using a pulled clear capillary glass tube (Harvard Apparatus 30-0062, 1.5 OD x 1.17 ID x 100mm). CSF was stored in a low binding microcentrifuge tube (Costar, 3206), then frozen on dry ice and stored at -80 degrees until assayed. After CSF collection, mice were transcardially perfused with 20ml 0.1M PBS. Brains were fixed in 4% PFA overnight at 4°C, before being transferred to 30% sucrose for cryoprotection. Brains were embedded in Tissue-Tek (Sakura) and sectioned coronally at 40µm using a sliding microtome (Microm HM450; Thermo Scientific).

#### Immunohistochemistry

For immunohistochemistry, a 1 in 6 series of 40µm coronal sections was incubated in 3% normal donkey serum in 0.3% Triton X-100 in TBS blocking solution at room temperature for one hour. Rabbit anti-IBA1 (1:1000, Wako Chemicals 019-19741), rat anti-CD68 (1:200, Abcam ab53444), rabbit anti-DCX (1:500, Abcam ab18723), rabbit anti-ASPA (1:250, EMD Millipore ABN1698), and goat anti-PDGFRa (1:500; R&D Systems AF1062) were diluted in 1% normal donkey serum in 0.3% Triton X-100 in TBS and incubated overnight at 4°C. Sections were then rinsed three times in 1X TBS and incubated in secondary antibody solution (Alexa 647 donkey anti-rabbit IgG, 1:500 (Jackson Immunoresearch); Alexa 488 donkey anti rat IgG, 1:500 (Jackson Immunoresearch); Alexa 594 donkey anti-rabbit IgG, 1:500 (Jackson Immunoresearch); Alexa 488 donkey anti goat IgG, 1:500 (Jackson Immunoresearch); Alexa 647 donkey anti-mouse IgG, 1:500 (Jackson Immunoresearch) in 1% blocking solution at room temperature protected from light for two hours. Sections were rinsed 3 times in TBS, incubated with DAPI for 5 minutes (1:1000; Thermo Fisher Scientific) and mounted with ProLong Gold mounting medium (Life Technologies).

#### Confocal imaging and quantification

All cell counting was performed by experimenters blinded to experimental conditions on a Zeiss LSM700 or LSM980 scanning confocal microscope (Zeiss). Images were taken at 20X magnification and analyzed using either FIJI software or Imaris software. For microglia imaging, three consecutive sections ranging approximately from bregma +1.2 to +0.8 were selected for cortex and corpus callosum, and four consecutive sections ranging approximately from bregma - 1.8 to -2.4 were selected for the hilus of the dentate gyrus. For each section, the superficial cortex, deep cortex, cingulum, genu of the corpus callosum, and hilus of the dentate gyrus were identified, and two 219.5 x 219.5 µm fields per slice were selected in those areas for quantification. All CD68^+^ and IBA1^+^ cells were counted in those regions. For analysis of immature neurons, four consecutive sections ranging approximately from bregma -1.8 to -2.4 were selected for the dentate gyrus, and four 219.5 x 219.5 µm fields per slice were selected in those areas for quantification. All DCX^+^ cells within the inner layer of the dentate gyrus were counted in those regions. For analysis of oligodendrocytes (ASPA^+^) and oligodendrocyte precursor cells (PDGFRa^+^) three consecutive sections ranging from bregma +1.2 to +0.8 were stained. The cingulum and genu of the corpus callosum in each hemisphere was imaged with two 319.5 x 319.5 µm field and cells were quantified using the “Spots” function in Imaris image analysis software (Oxford Instruments). Each image was manually inspected to ensure accurate oligodendrocyte counts. False positive and false negative counts were manually removed or added, respectively.

#### Electron Microscopy

At D35 post CAR T-cell treatment in the DIPG xenograft model, mice were anesthetized with isoflurane and CSF was collected as described above. Mice were then sacrificed by transcardial perfusion with Karnovsky’s fixative: 2% glutaraldehyde (Electron Microscopy Sciences (EMS 16000) and 4% paraformaldehyde (EMS 15700) in 0.1 M sodium cacodylate (EMS 12300), pH 7. Samples were prepared as detailed previously in^11^. Axons in the cingulum of the corpus callosum were analyzed for myelinated axon density (total number of myelinated axons in each frame) by experimenters who were blinded to experimental conditions and genotypes. A minimum of 500 axons were scored per group.

#### Mouse CSF and serum cytokine analysis

The Luminex multiplex assay was performed by the Human Immune Monitoring Center at Stanford University. Mouse 48-plex Procarta kits were purchased from Thermo Fisher, Santa Clara, California, USA, and used according to the manufacturer’s recommendations with modifications as described below. Briefly: beads were added to a 96-well plate and washed in a Biotek Elx405 washer. Samples were added to the plate containing the mixed antibody-linked beads and incubated overnight at 4°C with shaking. Cold (4°C) and room temperature incubation steps were performed on an orbital shaker at 500-600 RPM. Following the overnight incubation plates were washed in a Biotek Elx405 washer and then biotinylated detection antibody added for 60 minutes at room temperature with shaking. Plate was washed as described above and streptavidin-PE was added. After incubation for 30 minutes at room temperature, washing was performed as above and reading buffer was added to the wells. CSF samples were measured as singlets, while serum samples were measured in duplicate. Plates were read on a FM3D FlexMap instrument with a lower bound of 50 beads per sample per cytokine. Custom Assay Chex control beads were purchased from Radix Biosolutions, Georgetown, Texas, and were added to all wells.

#### Novel Object Recognition Task

Cognition was analyzed using a modified version of the novel object recognition task (NORT) that focused on the attentional component of the task. The test was modified so that the duration between the training and testing phase was shortened, to ensure that the test placed a greater cognitive load on short-term (< 5 min) memory, attention, and frontal lobe function rather than longer-term memory and hippocampal function. In DIPG, aggressive osteosarcoma and rapidly clearing osteosarcoma models, mice were tested via NORT on D34 post CAR-T cell administration. In the ALL model, mice were tested on D27 post CAR-T cell administration.

Animals were handled for 3 days leading up to the test for 2 min. After handling, the mice were then placed in the experimental chamber on D33 for the DIPG, aggressive osteosarcoma and rapidly clearing osteosarcoma models or D26 for ALL model to acclimatize for 20 min prior to testing on D34 or P27, respectively. The set up consisted of an opaque Plexiglas experimental chamber 50cm x 50cm x 50cm, and a camera mounted 115 cm above the chamber. The test was conducted during the animal’s rest phase, in a dark room illuminated with only red light.

On the day of testing, mice were brought to the testing room and allowed to settle in their cages for 20 minutes prior to any handling. Afterwards, mice were handled for 2 min and then placed in the chamber to acclimate for 20 min before being returned to the home cage for another 5 min. Mice were then placed in the experimental chamber with two identical Lego objects (each approximately 4 cm in size). Each time the mouse was placed into the chamber it was facing the opaque wall, with the animal’s tail in the direction of the objects. For the training phase, the mouse was allowed to explore the two identical objects for 5 min and then the mouse was returned to the home cage for 5 min while the experimental chamber and objects were cleaned with 1% Virkon S broad spectrum disinfectant. During this time, one cleaned object from the sample phase was placed back into the experimental chamber along with a new Lego object (of similar size) for the novel object phase. For the novel object testing phase, the mouse was returned to the experimental chamber and allowed to explore for 10 min.

The objects used as novel and familiar were counterbalanced, as was the position of the novel object from trial to trial, animal to animal. All of the Lego objects used in the behavioral paradigm were piloted to ensure that there was no bias, or object preference for the animals. The camera footage was then analyzed using Noldus analysis software and any exploratory head gesturing within 2 cm of the Lego object, including sniffing and biting, was considered object investigation, but sitting on the object, or casual touching of the object in passing was not considered object investigation (Leger et al., 2013). Only animals that explored both of the identical objects during the testing phase for an accumulation of a minimum of 20 s were included in the analysis. The Recognition Ratio was determined by taking the ratio of the amount of time spent investigating one object compared to the total time spent investigating both objects (i.e., time spent with Novel Object / (time spent with Novel Object + time spent with Familiar Object)).

#### Spontaneous Alternation T Maze Task

Hippocampal-dependent cognitive performance was analyzed using the spontaneous alternation T maze task, which focuses on assessing spatial working memory. This test was administered over 2 days beginning at the end of the NORT assessment. DIPG, aggressive osteosarcoma and rapidly clearing osteosarcoma models were tested on D34 and D35 post CAR-T cell administration. The ALL model was tested on D27 and D28 post CAR-T cell administration. At this point, mice had been handled prior to, and during NORT. The test was conducted during the animal’s rest phase, in a dark room illuminated with only red light. The T maze set up consisted of a start arm and two goal arms, each 30 X 10 cm with 20 cm high walls, where one goal arm of the T maze is textured to allow for clear distinction by the mouse^28^.

For day 1 of the T maze assessment, after NORT had been completed and mice given at least 2 hours of rest, a mouse is placed at the base of a T-shaped maze facing away from the goal arms. As the mouse explores the maze, it chooses an arm. After arm selection, by virtue of having four paws in that arm, a guillotine door was closed to allow the mouse to only explore and familiarize itself with the chosen arm for 30 s. The mouse was then placed back at the base of the maze facing away from the goal arms. This process was repeated for 11 consecutive cycles allowing for 10 total choices.

On day 2, mice were brought into the testing room and allowed to settle in their cages for 20 minutes prior to any handling. Afterwards, mice are each handled for 2 minutes prior to T maze testing. Next, a mouse was placed at the base of a T-shaped maze facing away from the goal arms. The mouse then chose an arm by having four paws in that arm. At that moment, the guillotine door was closed to allow the mouse to explore the chosen arm for 30 s. The mouse was then returned to its home cage for 15 min. The mouse was then placed back at the base of the maze facing away from the goal arms and allowed to explore and choose a goal arm. This process was repeated for 11 consecutive cycles. The chosen arms were recorded during testing and mice were scored over 10 trials.

### QUANTIFICATION AND STATISTICAL ANALYSIS

#### For all quantifications, experimenters were blinded to sample identity and condition

For fluorescent immunohistochemistry and confocal microscopy, images were taken at 20X and cells considered co-labeled when markers co-localized within the same plane. For cortical/white matter microglia staining three sections per mouse, eight frames in standardized locations per section (24 images per mouse) were counted, for hippocampal microglia staining, four sections per mouse, 2 frames in standardized location per section (8 images per mouse) were counted. For hippocampal neuroblast staining, four sections per mouse, 4 frames in standardized location per section (16 images per mouse) were counted. For oligodendrocyte staining three sections per mouse, 8 frames in standardized location per section (24 images per mouse) were counted. For each mouse, 300-700 cells were counted for each immunohistochemical marker analysis. The density of cells was determined by dividing the total number of cells quantified for each lineage by the total volume of the imaged frames (mm^3^). In all experiments, “n” refers to the number of mice. In every experiment, “n” equals at least 3 mice per group; the exact “n” for each experiment is defined in the figure legends.

For electron microscopy (EM) quantification of myelinated axon density, 6000X images were obtained. For myelinated axon density, the number of axons surrounded by a myelin sheath were counted and normalized to tissue area. 29 images were scored for each mouse. Approximately 400-700 axons were scored for each mouse.

All statistics were performed using Prism Software (Graphpad). For all analyses involving comparisons of one or two variables, 1-way or 2-way ANOVAs, respectively, were used with Tukey’s multiple comparisons post hoc corrections used to assess main group differences. For analyses involving only two groups, unpaired two-tailed Student’s t tests were used. For measurements of individual cytokine levels, unpaired two-tailed Student’s t tests were used without correction for multiple comparisons. Shapiro-Wilk test was used to determine normality for all datasets; all datasets were parametric. A level of p < 0.05 was used to designate significant differences.

